# Halogen-aromatic π interactions modulate inhibitor residence time

**DOI:** 10.1101/255513

**Authors:** Christina Heroven, Victoria Georgi, Gaurav K. Ganotra, Paul E. Brennan, Finn Wolfreys, Rebecca C. Wade, Amaury E. Fernández-Montalván, Apirat Chaikuad, Stefan Knapp

**Affiliations:** Nuffield Department of Clinical Medicine, Structural Genomics Consortium, University of Oxford, Oxford, OX3 7DQ, UK; Bayer AG, Drug Discovery, Pharmaceuticals, Lead Discovery Berlin, 13353 Berlin, Germany; Buchmann Institute for Molecular Life Sciences, Johann Wolfgang Goethe-University, D-60438 Frankfurt am Main, DE; Institute for Pharmaceutical Chemistry, Johann Wolfgang Goethe-University, D-60438 Frankfurt am Main, DE; Molecular and Cellular Modeling Group, Heidelberg Institute for Theoretical Studies (HITS), 69118 Heidelberg, Germany; Target Discovery Institute, Nuffield Department of Clinical Medicine, University of Oxford, Oxford, OX3 7FZ, UK; German Cancer Network (DKTK), Frankfurt/Mainz site, D-60438 Frankfurt am Main, DE; Zentrum für Molekulare Biologie, DKFZ-ZMBH Alliance, Heidelberg University, 69120 Heidelberg, Germany; Interdisciplinary Center for Scientific Computing, Heidelberg University, 69120 Heidelberg, Germany

**Keywords:** Halogen-π interaction, drug residence time, kinase

## Abstract

Prolonged drug residence times may result in longer lasting drug efficacy, improved pharmacodynamic properties and “kinetic selectivity” over off-targets with fast drug dissociation rates. However, few strategies have been elaborated to rationally modulate drug residence time and thereby to integrate this key property into the drug development process. Here, we show that the interaction between a halogen moiety on an inhibitor and an aromatic residue in the target protein can significantly increase inhibitor residence time. By using the interaction of the serine/threonine kinase haspin with 5-iodotubercidin (5-iTU) derivatives as a model for an archetypal active state (type I) kinase-inhibitor binding mode, we demonstrate that inhibitor residence times markedly increase with the size and polarizability of the halogen atom. This key interaction is dependent on the interactions with an aromatic residue in the gate keeper position and we observe this interaction in other kinases with an aromatic gate keeper residue. We provide a detailed mechanistic characterization of the halogen-aromatic π interactions in the haspin-inhibitor complexes by means of kinetic, thermodynamic, and structural measurements along with binding energy calculations. Since halogens are frequently used in drugs and aromatic residues are often present in the binding sites of proteins, our results provide a compelling rationale for introducing aromatic-halogen interactions to prolong drug-target residence times.

## Introduction

The kinetics of drug binding have emerged as important parameters in drug development. A long drug residence time will result in prolonged inhibition after the free drug concentration has dropped due to *in vivo* clearance, potentially leading to improved drug efficiency and reduced off-target mediated toxicity. In cases where slow off-rates are specific for the target, kinetic selectivity can be achieved over fast off-target dissociation despite similar binding constants(1). The kinetics of the interaction of a drug with its target are defined by the association rate (k_on_) and the dissociation rate (k_off_) constants. For bimolecular interactions, the ratio of these two parameters defines the equilibrium dissociation constant (K_D_) of a drug, and hence the drug occupancy. Since on‐ and off-rates are coupled in simple rigid bimolecular interactions, the favorable scenario of fast “on” and slow “off”-rate interactions at a given K_D_ cannot be achieved using this minimalistic binding model. Instead more complex binding models such as induced conformational changes that may trap a ligand in an induced binding pocket are frequently evoked to explain slow binding kinetics.

Kinases are particularly dynamic proteins that provide multiple opportunities for the development of inhibitors that target induced or allosteric binding sites (2-4). One of the first kinase inhibitors for which slow dissociation rates have been described is the p38 MAP (<u>m</u>itogen <u>a</u>ctivating <u>p</u>rotein) kinase inhibitor BIRB-796. This inhibitor binds to an inactive conformation in which the DFG motif is displaced in a so-called “DFG-out” conformation (5). Inhibitors that bind to this conformation are called type II inhibitors and often have prolonged residence times (τ). However, not all type II inhibitors show slow binding kinetics, suggesting that the DFG-out conformational change *per se* is not sufficient to explain the slow dissociation rates of BIRB-796 from p38 MAP kinase(6). Indeed, more recent studies attributed the slow binding kinetics to efficient hydrophobic contacts in the DFG-out pocket, rather than the kinetic dissociation barrier introduced by the DFG-out transition(7). However, conformational change has also been evoked as the main mechanism contributing to the slow off-rate of the breast cancer drug lapatinib, a type I inhibitor of the epidermal growth factor receptor (8).

In addition to protein conformational changes, the rearrangement of water molecules has been discussed as a potential mechanism influencing inhibitor residence time (9). An example for the influence of water molecules on ligand binding kinetics is the type I CDK inhibitor roniciclib whose slow off-rate is the result of changes in the hydration network coupled to conformational adaptation of the DFG motif (10). In some cases, the presence of water-shielded hydrogen bonds can also lead to slow dissociation behavior (11). In addition, reversible covalent inhibitors have recently emerged as an interesting strategy for prolonging target engagement of inhibitors by the interaction of a transient covalent bond between an electrophile and a cysteine residue present in the kinase active site (12-15).

Even though protein conformational changes and allosteric pockets can be specifically targeted, they do not offer a straightforward route for ligand design for the modulation of off-rates. Also, the design of reversible covalent interactions requires the presence of cysteine residues in the drug binding site. Many drug receptors, including a large number of kinases, contain cysteines in close proximity to their active sites (16), but the development of covalent inhibitors may not be feasible for all drug targets. In addition, the introduction of a reactive group into an inhibitor poses additional challenges.

Here, we present data that suggest that interactions mediated by halogens, that are common in drugs, and aromatic residues, that are also typically found in drug binding sites on proteins (17), can be utilized to design ligands with slow off-rates. We used 5-iodotubercidin (5-iTU), a close analogue of the kinase cofactor ATP, as a model inhibitor for a canonical active state kinase binding mode (type I) which does not induced any conformational changes that could contribute to slow inhibitor binding kinetics. Screening against more than 100 diverse kinases showed that an aromatic gatekeeper residue that interacts with the halogen moiety of this inhibitor is required for high affinity binding. We chose haspin, a serine/threonine kinase with known three-dimensional structure (18, 19), as a model system. Analysis of ligand binding kinetics surprisingly showed that 5-iTU had slow binding kinetics. Mutation of the gatekeeper residue as well as removal or substitution of the iodide with other halogens showed that the slow inhibitor off-rates are due to a π-stacking interaction of the halogen with the aromatic gatekeeper. Further studies on a different kinase, cdc2 like kinase (CLK1), showed strong inhibition by 5-iTU, support the generality of our findings. Here, we present structural, biophysical and computational data that strongly suggest that halogen interactions with aromatic residues can be exploited for the development of inhibitors with slow off-rates.

## Results and Discussion

### 5-iTU exhibited slow dissociation rate from haspin

Comparative analysis of the high resolution haspin structures revealed high conservation of the binding modes of both 5-iTU and the nucleoside adenosine (Fig. 1A-B). In contrast to adenosine or ATP, which binds with a K_D_ of about 180 μM(19), 5-iTU showed high affinity for haspin and an unexpectedly long target engagement time. We then further assessed the thermodynamics and kinetics of the binding with ITC (Isothermal Titration Calorimetry), BLI (BioLayer Interferometry) and SPR (Surface Plasmon Resonance) experiments, which consistently confirmed the tight binding with a relatively slow binding kinetics, as measured for instance by BLI (Fig. 1C). Comparing the binding mode of 5-iTU with that of adenosine, the most striking structural difference between these highly similar molecules was the presence of the iodide moiety which was positioned in close proximity to the F605 gatekeeper forming a halogen-aromatic π interaction (Fig. 1D). We therefore hypothesized that this interaction might contribute most of the increase in binding free energy (ΔG) and be responsible for the slow dissociation rate of 5-iTU from haspin.

**FIGURE 1.**
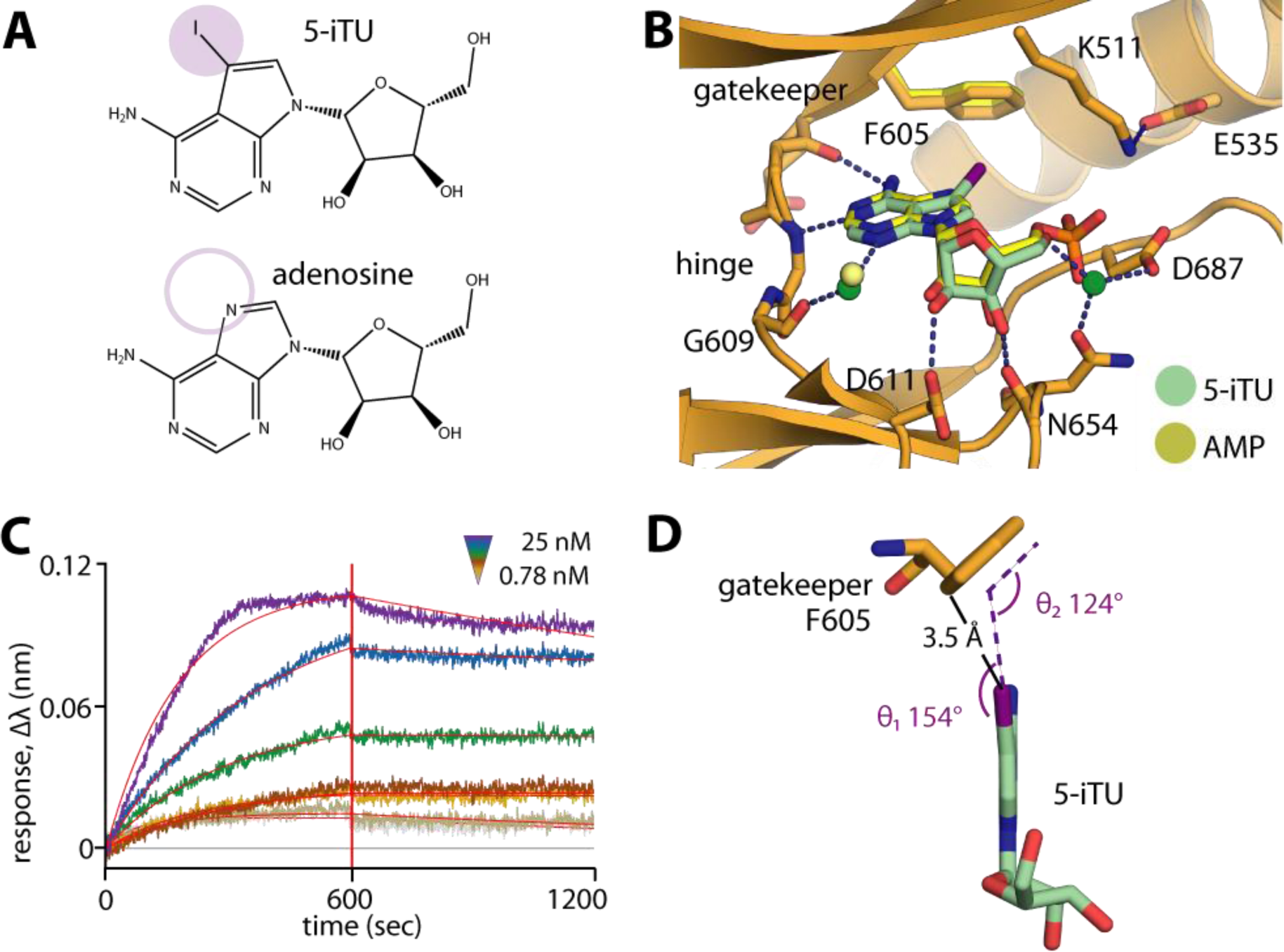
The 5-iodotubercidin inhibitor (5-iTU) exhibits tight binding with slow dissociation kinetics from haspin. A) Chemical structures of 5-iTU and adenosine. B) Superimposition of haspin-5iTU and AMP (pdb id: 4ouc) reveals similar binding modes of the two compounds. C) BLI sensorgram suggests slow kinetic behavior of the 5-iTU-haspin interaction. D) The iodide and the benzene moieties of 5-iTU and F605, respectively, are located in close proximity with a favorable geometry for a halogen-π bond.

### Preference of 5-iTU for kinases with aromatic gatekeeper

To address our hypothesis, we screened 5-iTU against 137 diverse kinases using temperature shift assays (20) and observed unexpected selectivity of the inhibitor with only 10 kinases exhibiting ΔT_m_ of >5 °C (Fig. 2A, Table S1). Interestingly, analyses of the gate-keeper residues of kinase targets that showed significant temperature shifts and therefore high affinity for 5-iTU, revealed a strong preference for kinases harboring a phenylalanine (Phe) at this position whereas kinases that showed weak interaction (ΔT_m_ 2-5 °C) revealed no preference for a certain residue. This analysis supported our hypothesis that the aromatic gatekeeper is important for high affinity binding (Fig. 2B). To confirm these results, we determined the structure of 5-iTU bound to another high affinity target that is structurally very diverse from haspin, CLK1 (ΔT_m_ of 8.6 °C). As expected, the interaction of 5-iTU with CLK1 remarkably resembled that observed in haspin, including the conserved interaction geometry of the iodide with the CLK1 gatekeeper F241. 5-iTU bound CLK1 with high affinity (K_d_ of ~7 nM by ITC) and slow off-rates estimated to be ~50 mins by BLI (Fig. 2C-E and Fig. S1).

**FIGURE 2.**
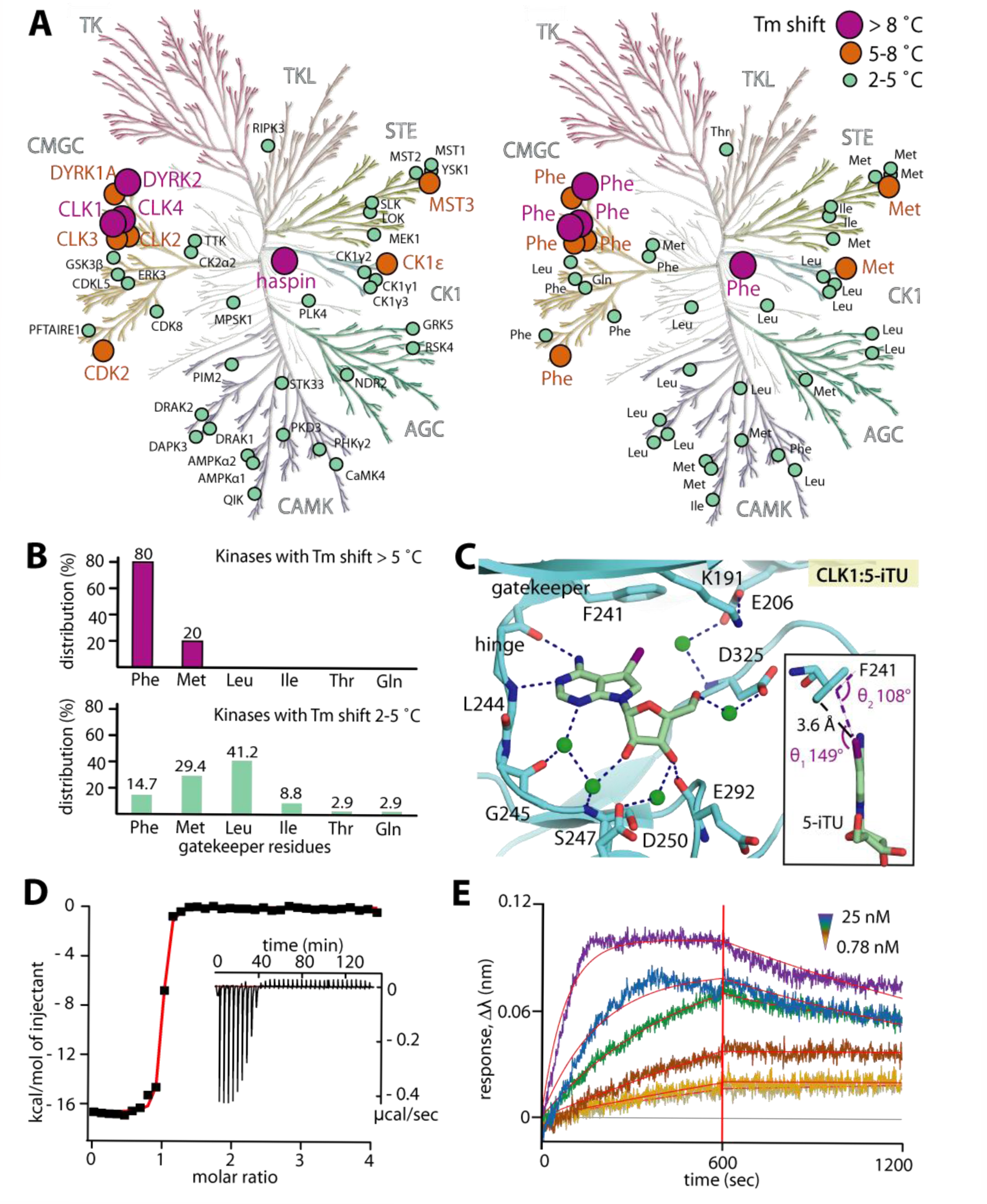
Interaction of 5-iTU with various human kinases. A) Temperature shift assays show high ΔT_m_ preferentially for kinases harboring an aromatic gatekeeper. B) Distribution of gatekeeper residues reveals that the majority of strong ΔT_m_ hits harbor a phenylalanine gatekeeper. C) The binding mode of 5-iTU in the off-target CLK1, including the iodide gatekeeper interaction, is highly conserved. CLK1 has a high affinity for 5-iTU as measured by ITC (D) and a slow off-rate (E) as assessed by BLI.

### Importance of halogen σ-hole property for halogen-π interaction for slow off-rate

Formation of a halogen-π interaction (C-X···π) is driven by the directional positive polarization along the halogen σ-bond with the π molecular orbital of the aromatic system (21). However, the main focus of halogen-protein interactions has been on halogen carbonyl/sulphonyl interaction (C-X···O,S bonds) with few examples on the analysis of halogen-aromatic interactions (22). Because the partial positive charge along the halogen σ-bond diminishes with the size of the halogen, we next substituted the iodide by smaller halogens and characterized the affinities and binding kinetics of these 5-iTU derivatives. Indeed, ITC experiments showed that the affinities of 5-iTU halogen derivatives were reduced with decreasing size of the halogen (Fig. 3 and Table S2).

**FIGURE 3.**
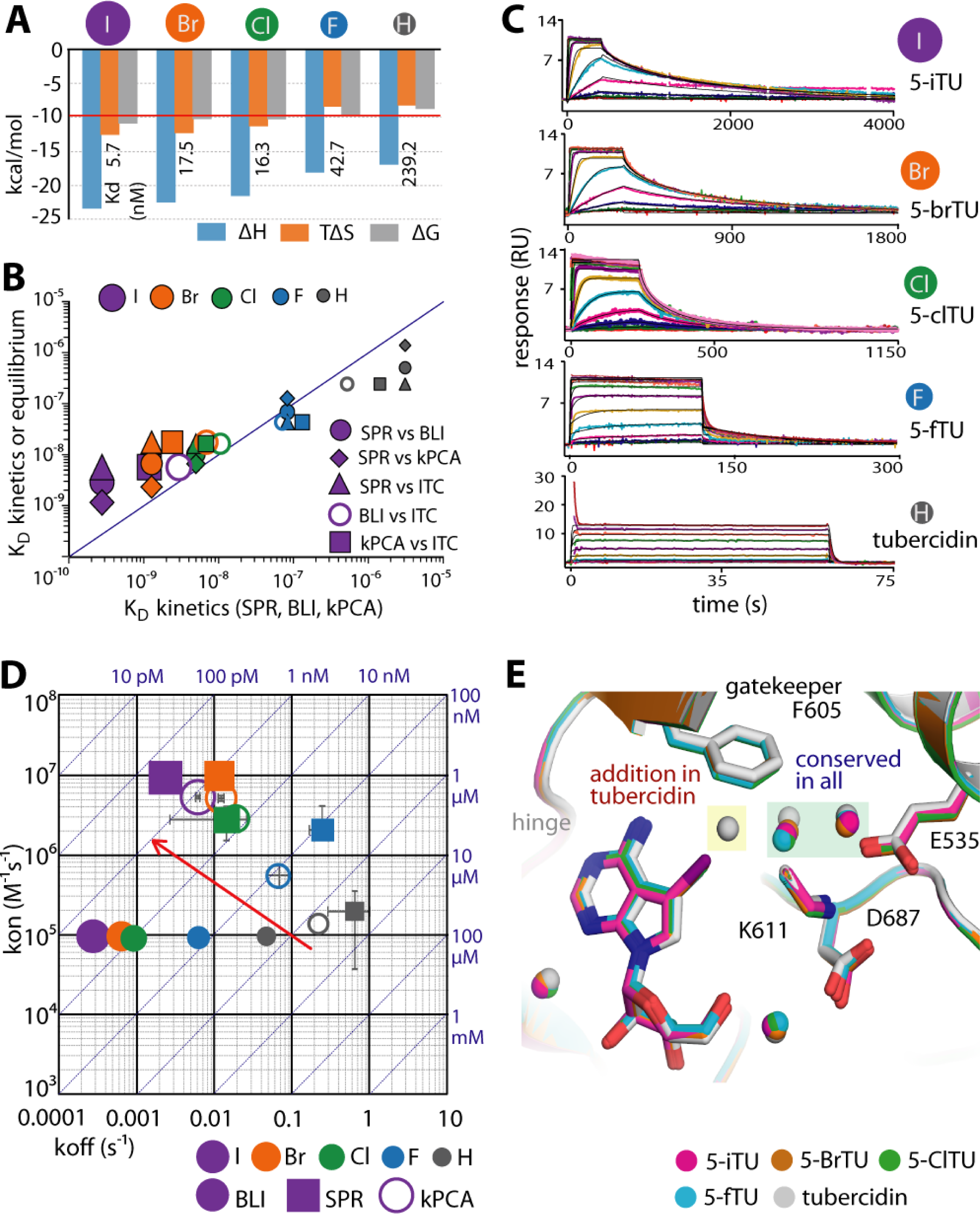
Binding kinetics of haspin with five tubercidin derivatives harboring halogen substituents at the 5-position. A) ITC thermodynamic binding parameters. B) Comparison of dissociation constants (K_D_) measured by ITC, BLI, SPR and kPCA shows good correlation of the measured equilibrium data. C) SPR sensorgrams demonstrating increasingly slow dissociation rates with increasing size of the halogens. D) Rate plot with Isoaffinity Diagonal (RaPID) of k_on_ and k_off_ constants measured by BLI, SPR and kPCA. The red arrow indicates the trend to increasing k_on_ and decreasing k_off_ upon increasing the atomic radii of the halogens. E) Crystal structures reveal conserved binding modes of all five tubercidin derivatives, albeit with an additional water molecule adjacent to the inhibitor and F605 gatekeeper in tubercidin.

Removal of the halogen led to a 42-fold decrease in the potency of tubercidin (TU) when compared to that of 5-iTU, and similarly an 8-fold decrease was measured for 5-fluorotubercidin (5-fTU). Analyses of the binding kinetics of these five synthesized 5-tubercidin halogen derivatives with haspin were performed using three independent techniques: kinetic probe competition assays (kPCA) in solution(23), BLI and SPR using immobilized haspin. Binding affinities determined by these three independent methods correlated well with each other and also with the binding constants determined in solution using ITC (Fig. 3B). Dissociation rate constants from all experiments revealed the same behavior with 5-iodide substituted tubercidin displaying the slowest off-rate. The off-rates increased with decreasing halogen size from 5-iodo to 5-fluoro substituted tubercidin and the unsubstituted tubercidin showed the fastest off-rates of binding (Fig. 3C-D). However, the absolute values differed somewhat between the different experimental methods used with the residence time ranging from 60 mins (BLI) to 7 mins (SPR) for 5-iTU (SI Appendix, Table S2 and S3). We also observed slower on rates using BLI but not in the SPR experiments. While the general trends were the same in both technologies, the differences in on‐ and off-rates that have been observed might be due to differences in protein immobilization. As SPR is the more established technology, we used SPR kinetic data for quantitative analysis. The substitution from hydrogen to iodide at the 5 position of TU led to a 48-fold increase in the on-rates and 274-fold decrease in the off-rates (Fig. 3D and SI Appendix, Table S3).

Thus, the fast dissociation kinetics observed for 5-fTU coincided with the lack of a pronounced σ-hole in smaller halogens, and hence an inability to form a polar halogen-π interaction with the aromatic gatekeeper. The increase in enthalpically favorable polar interactions of larger halogens was also evident in the calorimetric data that showed a steadily decreasing (more negative) binding enthalpy change from tubercidin to the larger halogens (H < F < Cl < Br < I). This effect of the halogen moieties was supported by the crystal structures showing that despite the highly conserved binding mode of all derivatives, the presence of an additional water in the tubercidin complex within the space adjacent to the gatekeeper created by the removal of larger halogen substituents and the longer distance to the fluoro group, presented suboptimal geometry for direct contacts with F605 in tubercidin and 5-fTU, respectively (Fig. 3E and SI Appendix, Fig. S2).

### Importance of aromatic phenylalanine for slow off-rate binding

We next investigated the contributions of the Phe gatekeeper to the binding affinity and the observed slow dissociation kinetics of 5-iTU. Six haspin gatekeeper mutants using amino acids commonly found in kinases were generated, and their affinities against 5-iTU and its halogen-derivatives were analyzed using DSF assays (Fig. 4A). As expected, the potency of 5-iTU with the aromatic tyrosine mutant was comparable to wild-type haspin.

**FIGURE 4.**
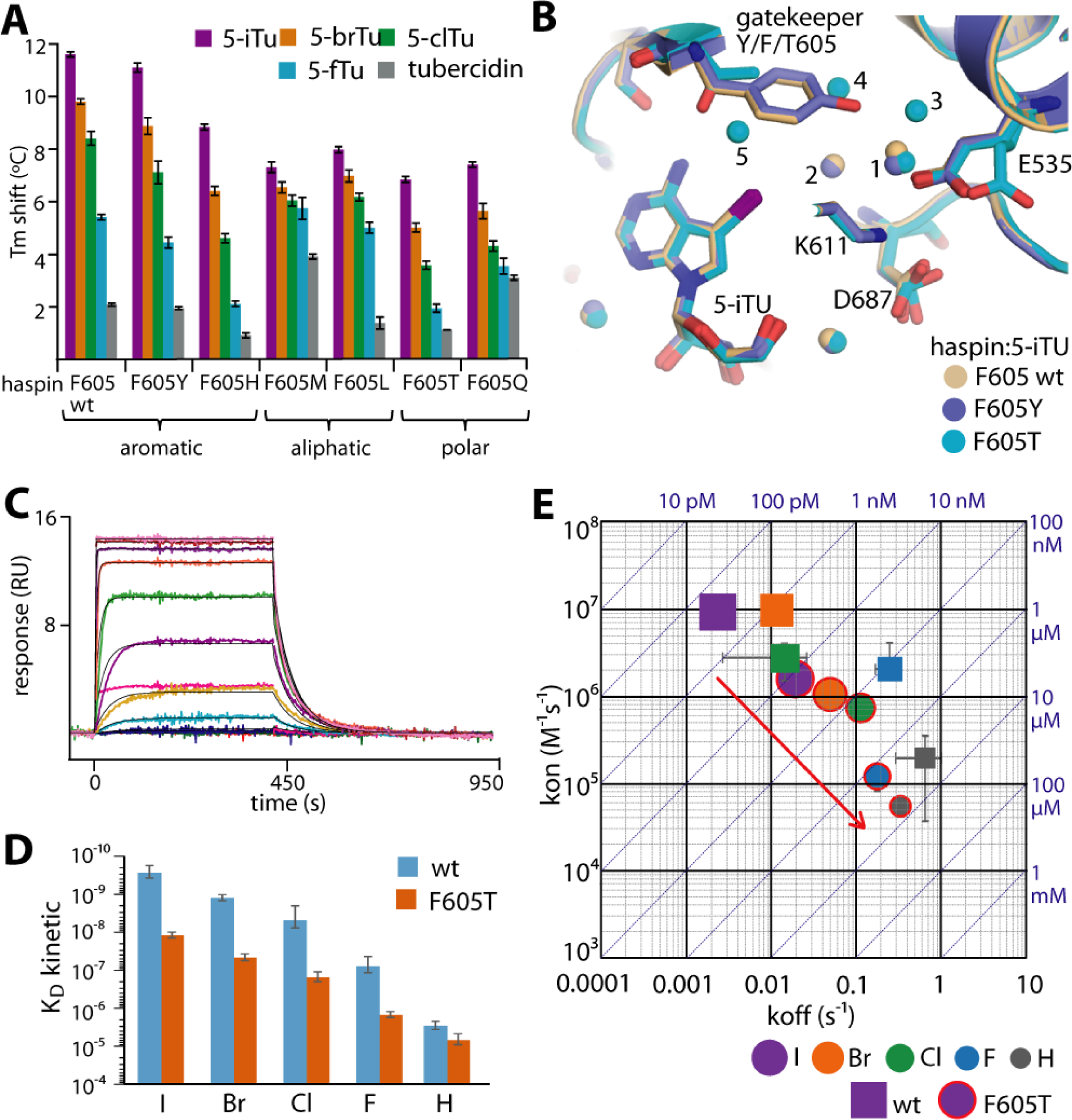
Effect of gatekeeper mutation on the binding kinetics of haspin with tubercidin derivatives. A) Tm shifts of six haspin mutants against five tubercidin derivatives. B) Superimposition of 5-iTU-complexed crystal structures of wild-type, F605Y and F605T haspin reveals conserved binding mode of the inhibitor, yet differences in bound water molecules within the binding site. C) SPR sensorgram demonstrates fast binding kinetics for the interaction between 5-iTU and the F605T mutant, and this is accompanied by a significant decrease in K_D_ (D) and decreased k_on_ and generally increased k_off_ constants (E).

For comparison, we analyzed the binding characteristics of 5-iTU with a representative mutant (F605T) in detail. Structural superimposition between the wild-type and the mutant structures (F605Y and F605T) showed that the gatekeeper mutation did not affect the binding mode of 5-iTU, yet led to slight variation of the environment in the pocket. The substitution of the bulky aromatic residue F605 with the small threonine resulted in extension of a water network that filled the expanded binding site in the mutant (Fig. 4B and SI Appendix, Fig. S3). No direct contact was observed between the threonine side chain and the iodide of 5-iTU, although interactions might be mediated through a water bridge. The absence of any strong contact was in agreement with the fast kinetics with ~16-fold lower affinity as demonstrated by SPR (Fig. 4C-D). In comparison to the wild-type, the loss of the halogen-π contact in the F605T mutant led to a 6-fold decrease and 8-fold increase in association and dissociation rates, respectively, with the estimated residence time of 5-iTU dramatically dropping to less than a minute (Fig. 4E and SI Appendix, Table S3).

### Thermodynamics of halogen-π interactions in slow kinetic behavior

In order to assess the energetic contributions of the halogen-aromatic gatekeeper interaction, we calculated the energy of this interaction using *ab initio* quantum mechanics and classical methods. The second order Møller–Plesset interaction energies (E_MP2_) between the inhibitor and the gatekeeper residue correlate well with dissociation rate constants and equilibrium dissociation constants determined experimentally (Fig. 5A and SI Appendix, Fig. S5 and Tables S6-S9). Partitioning of E_MP2_ into its constituent energy components using a many-body interaction energy decomposition scheme shows that the major contribution to E_MP2_ is the correlation energy (E_CORR_) which describes second-order intermolecular dispersion interactions and the correlation corrections to the Hartree-Fock energy. E_CORR_ increases in magnitude with increasing size of the halogen, corresponding to the decreasing rate of dissociation measured experimentally (Fig. 5B), and indicating the importance of the halogen interaction with the aromatic gatekeeper for the prolongation of residence times as halogen size increases. The computed *ab initio* energies also correlate for the interaction of 5-iTU with the F605Y mutant but the magnitude of the interaction energy of 5-iTU with the threonine mutant was underestimated. This discrepancy could be explained by the additional bound water molecule present between 5-iTU and the threonine residue in the F605T structure which was not included in the calculation but would make an energetically favorable contribution to the binding. To account for the complete protein structure in the computation of the binding free energies of the haspin-ligand complexes, we used the classical MMGBSA approach with an implicit solvent model. The computed energies correlate well with calorimetric data measured by ITC, consistent with the increasingly favorable enthalpic contribution to binding as halogen size increases (Fig. 5C-D and SI Appendix, Table S10).

**FIGURE 5.**
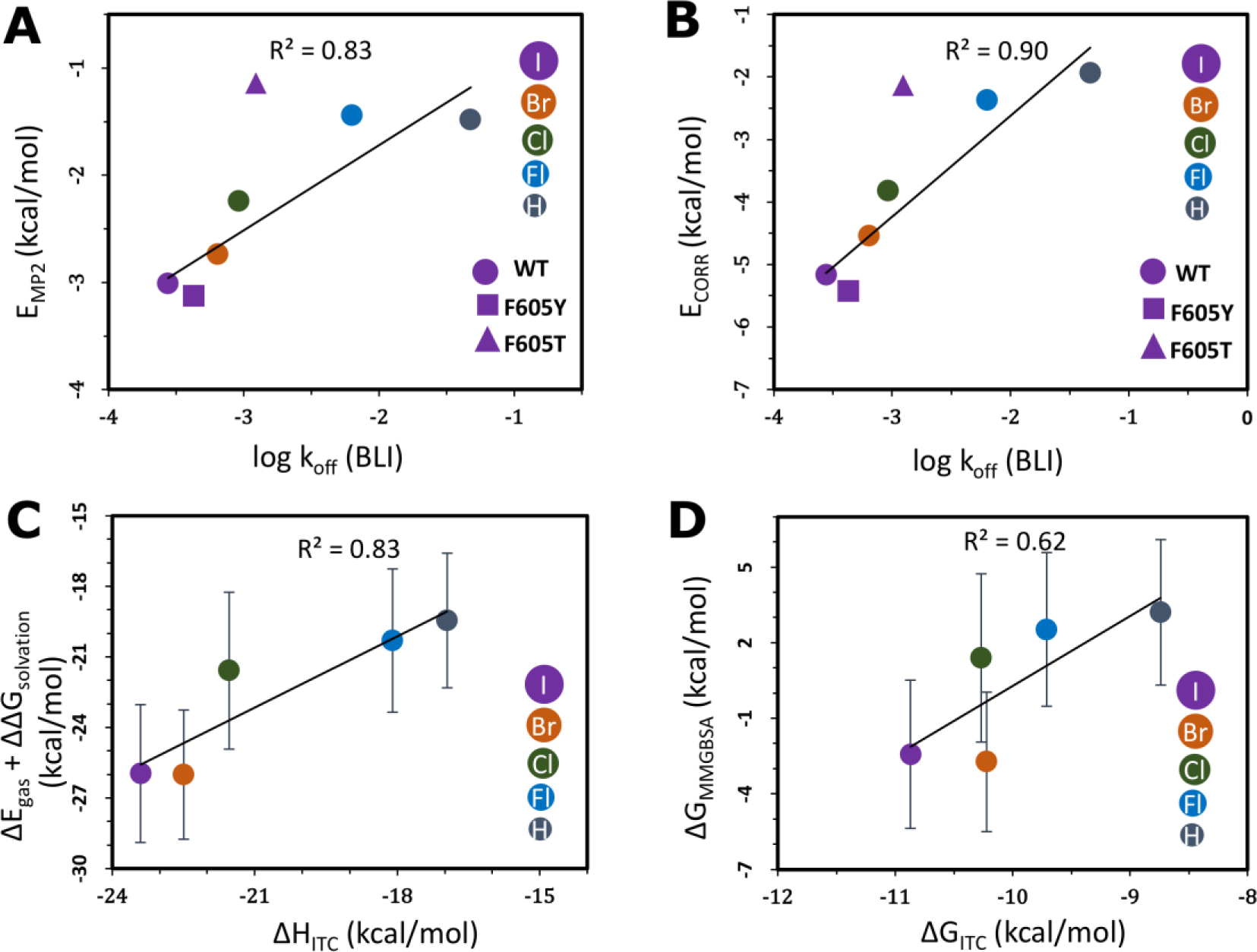
Correlation of calculated binding free energies with experimental parameters for the halogen-gatekeeper interaction. A) Second-order Møller–Plesset interaction energy (E_MP2_) and B) second-order correlation correction energy term (E_CORR_) between the TU derivatives and the gatekeeper residues *vs* BLI k_off_ values. The linear fits and correlation coefficients (R^2^) were computed omitting the outlier F605T mutant. The experimental error bars are smaller than the size of the data plots. Comparison of (C) MMGBSA internal and solvation contributions (ΔE_gas_+ΔΔG_solvation_) *vs* the ITC enthalpies (ΔH_ITC_) and (D) MMGBSA binding free energies (ΔG_MMGBSA_) *vs* the ITC binding free energies (ΔG_ITC_) of the interactions between haspin and TU derivatives. Some ΔG_MMGBSA_ values are positive as they only include translational and rotational entropic terms and do not include vibrational and conformational entropy contributions.

Halogens are frequently found in approved drugs. Interestingly, drug candidates with heavy halogens (Cl, Br, I) perform better in the clinical development pipeline increasing steadily from phase II (44%) to approved drugs (63%) whereas fluorine-containing drug candidates decrease from 56% in phase II to 36.6% in launched drugs (17). Unfortunately, information on binding kinetics is not available for these molecules but it is tempting to speculate that the presence of halogens may lead to longer lasting drug-protein interactions. A recent survey of the PDB for halogen-protein interactions reported that 33% of all non-bonded interactions (excluding C-X···H contacts) of the heavier halogens in the protein database form aromatic stacking interactions of the C-X⋯π type (24). A large number of halogens also interact with backbone carbonyls and thiols and it would be interesting to investigate if these interactions also result in slower ligand binding kinetics. For the selected case of protein kinases, aromatic gatekeeper residues are the most frequently found residue type. Our proposed strategy of incorporating heavier halogens into inhibitors with the goal of increasing target residence time by designing interactions with aromatic residues is a general approach that may lead to active compounds with improved pharmacological properties. The biophysical, structural and computational data presented here on 5-halogen substituted tubercidin derivatives provides a good basis for future studies on this exciting topic.

## ACKNOWLEDGMENT

This study was supported by the European Union’s Seventh Framework Program (FP7/2007-2013) for the Innovative Medicine Initiative under grant agreement no. 115366. A.C. is supported by the SFB/DFG program autophagy. G.K.G. and R.C.W. acknowledge the financial support of the Klaus Tschira Foundation. P.B. A.C. and S.K. are grateful for support by the SGC, a registered charity (no. 1097737) that receives funds from AbbVie, Bayer Pharma AG, Boehringer Ingelheim, Canada Foundation for Innovation, Eshelman Institute for Innovation, Genome Canada through Ontario Genomics Institute [OGI-055], Innovative Medicines Initiative (EU/EFPIA) [ULTRA-DD grant no. 115766], Janssen, Merck KGaA, MSD, Novartis Pharma AG, Ontario Ministry of Research, Innovation and Science (MRIS), Pfizer, São Paulo Research Foundation-FAPESP, Takeda, and the Wellcome Trust. We thank Diamond Light Source for support.

## Author Contributions

C.H., V.G, G.K.G., F.W. performed experiments, R.C.W., A.E.F-M., P.B., A.C., S.K supervised the research, S.K., A.C. drafted the manuscript with contributions from all authors. All authors approved the final version.

## Notes

The authors declare no competing financial interests.

## Material and Methods

### Protein purification

Haspin and CLK1, both wild-type and all mutants, have been purified as described (18, 25). Briefly, the recombinant proteins were purified using Co^2+^/Ni^2+^ affinity chromatography. For haspin, the His-tagged protein were subsequently purified by size-exclusion chromatography and the final protein was stored in 50 mM HEPES, pH 7.5, 300 mM NaCl, 0.5 mM TCEP. For CLK1, the histidine tag was removed by incubating the protein with TEV protease overnight. The cleaved CLK1 protein was separated by reverse purification on Ni^2+^ affinity chromatography, and subsequently size exclusion chromatography. The final CLK1 protein was stored in 30 mM HEPES, pH 7.5, 300 mM NaCl, 50 mM L-Arginine/L-Glutamate mix, 10 mM DTT, 1% glycerol.

### Protein crystallization

All crystallization experiments were performed using sitting-drop vapour-diffusion method at 4 °C. For Haspin, the protein at ~12 mg/ml was incubated with 1 mM inhibitors, and the complexed crystals were obtained using the crystallization condition containing 51-63% MPD and 0.1M SPG buffer, pH 6.0-6.5. To obtain the inhibitor-CLK1 complex, apo crystals grew in 20% 1,2-propanediol, 5% glycerol and 0.1 M NaKPO_4_ were soaked with inhibitor overnight.

### Data collection and structure determination

All diffraction data were collected at Diamond Light Source, and processed using MOSFLM(26). Scaling was performed using aimless from the CCP4 suite(27). All structures were solved by molecular replacement using Phaser (28) and the deposited CLK1 and haspin structures (PDB entries XYZ and 4OUC respectively) as models. All structures were subjected to one round of automated model building using *ARP/wARP(29)*, followed by iterative cycles of manual model building in COOT (30), alternated with refinement using REFMAC (31). TLS definitions used in the final refining rounds were calculated using the TLSMD server (31). The model quality and geometric correctness of all complexes was verified using MolProbity (32). Statistics for data collection and structure refinement are summarized in Supplemental Table 11.

### Thermal shift assays

Protein were diluted to 2 μM in a buffer containing 10 mM HEPES, pH 7.5 and 500 mM NaCl, and mixed with SYPRO Orange at 1000-fold dilution of the dye. The inhibitors were added at 10 μM final concentration. The DSF assay was performed using a Real-Time PCR Mx3005p machine (Stratagene) according to the protocol described previously (33).

### Isothermal Titration Calorimetry

All proteins were exchanged into a suitable storage buffer. CLK1 at 100 μM was stored in 20 mM HEPES, pH 7.5, 300 mM NaCl, 50 mM L-arginine/L-glutamate mix and 0.5 mM TCEP. For haspin at 80 μM, the buffer containing 20 mM HEPES, pH 7.5, 250 mM NaCl and 0.5 mM TCEP was used for the wild-type protein, while the gatekeeper mutants were buffer exchanged into 30 mM HEPES, pH 7.5, 400 mM NaCl and 0.5 mM TCEP to increase their stabilities. Calorimetric measurements were carried out using a VP-ITC calorimeter (MicroCal) at 15 °C. For all experiments, the proteins were titrated into the reaction cell containing the compound. Integrated heat of titrations were manually corrected and analysed in Origin. Using a single binding site model, the obtained curve was fitted following a nonlinear least-square minimization algorithm. The binding isotherms and the measured binding enthalpy changes enabled the calculation of entropy changes (TΔS), Gibbs free energy (ΔG), the stoichiometry *n* and K_d_.

### Biolayer Interference

The binding kinetics were measured by Biolayer Interference method (BLI) using Octet RED384 system (*forté*BIO). For haspin, the experiments were performed in the buffer condition containing 20 mM HEPES, pH 7.5, 400 mM NaCl and 0.5 mM TCEP. For CLK1, the same buffer supplemented with 50 mM L-arginine/L-glutamate mix was used. Biotinylated proteins, prepared as previously described, were immobilized on streptavidin biosensors, which were subsequently quenched with L-biotin(34). The interference patterns of association and dissociation events were measured through a time course of 600 seconds. The binding data were corrected using a double-referencing method, and the kinetics analyses were performed according to the manufacture protocol (*forté*BIO).

### Surface Plasmon Resonance (SPR)

SPR experiments were performed in HBS-PE+ buffer on a Biacore T200 System (GE Healthcare). Biotinlyated wt and mutant Haspin (50 μg/ml in HBS-P+ buffer) were captured to a SA sensor chip (GE Healthcare) at typical densities of 2-5 kRU using the protocols provided by the manufacturer. Compounds were serially diluted in DMSO and transferred to assay buffer in a 1:100 dilution step to achieve their final test concentrations at a [DMSO] = 1%. For binding analysis, contact times of 60, 120, 240, 300 or 420 seconds were used depending on the kinetics assessed in preliminary tests. Likewise, dissociation times were adjusted to180, 900, 1500 or 3600 seconds, to achieve return of the SPR signals to baseline levels. To obtain kinetic and affinity parameters, sensorgrams (acquired at 10 Hz) were fitted using the BIAevaluation Software (GE Healthcare) to a 1:1 Langmuir model accounting for mass transport limitations. Steady state analysis was performed with the same software using a single site equilibrium binding equation.

### Equilibrium und Kinetic probe competition assays (ePCA and kPCA)

ePCA and kPCA experiments were performed in Tris-HCl pH7.5, 150 mM NaCl, 0.01% Tween, 0.01% BSA, 2 mM DTT buffer as previously described for CDK2 in Schiele et al (23). Biotinylated wt and mutant Haspin (4 nM in assay) were labelled at a molar ratio of 8:1 with SA-Terbium (Cisbio) as TR-FRET donor. Tracers 236 and 199 (Invitrogen) labelled with an Alexa 647 TR-FRET acceptor were respectively used as kinase specific probes at a final concentration of 100 nM.

Compounds were diluted and transferred to Greiner black small volume 384-well microtiter test plates as described(23). For ePCA, tracer and labeled proteins were dispensed to the ready-to-use compound plates to a final volume of 5 μL and the mixture was incubated for 2 h prior to acquisition of the steady state TR-FRET ratiometric signals (665/620 nm) upon excitation at 337 nm. Normalized values were fitted to a logistic 4-parameter model using the Genedata Screener^TM^ software, and Ki values calculated using the Cheng-Prusoff relationship. For kPCA, the tracer was dispensed to the ready-to-use compound plates prior to introducing them into the PHERAstar FS^TM^ microtiter plate reader. Then the labeled proteins were added to wells to a final volume of 10 μL using the injector system of the instrument, and kinetic TR-FRET readings were made at time zero and every 10 seconds. Blank-subtracted kinetic traces were analyzed with a competitive binding kinetics model using the GraphPad Prism^TM^ software as described (23).

Prior to compound testing, the steady state affinities of the probes were determined by equilibrium binding titrations (0 to 400 nM) on various Haspin concentrations (0 to 8 nM) with end-point readings of the TR-FRET signals. The probes’ association and dissociation kinetics were characterized by titrating them on 4 nM labeled Haspin (0.5 nM SA-Tb) and acquiring the TR-FRET signals in real time. Binding curves were fitted to the corresponding models with Graph Pad Prism^TM^ in order to obtain the affinity and kinetic constants used as parameters in the Cheng-Prusoff and Motulsky and Mahan models.

### Quantum mechanical interaction energy calculations

The energy contributions of the inhibitor-aromatic gatekeeper interaction were calculated using *ab initio* Møller–Plesset perturbation theory to second order (MP2). The Moeller-Plesset perturbation theory improves on the Hartree-Fock method by adding electron-correlation effects by means of Rayleigh-Schrödinger perturbation theory to different orders (second order in our case). The *Protein Preparation* wizard of the *Maestro* program of the *Schrodinger suite* (*Version 2015.r3*) was used to pre-process the X-ray crystallographic structures of the haspin-inhibitor complexes, to add missing side chains and to optimize the H-bond network. The *impref* utility of the *Maestro* was used for energy minimization using the *OPLS3* force field. The *impref* utility(35) first optimizes position of hydrogen atoms followed by all-atom minimization where non-hydrogen atoms are restrained with a harmonic potential using a force constant of 25 kcal/mol.Å^2^. The coordinates of the inhibitor and the gatekeeper phenylalanine residue were extracted from these energy-minimized structures of haspin-inhibitor complexes. The termini of the phenylalanine residue were blocked with hydrogen atoms and their positions were optimized using the *OPLS3* force field in the *Maestro* program of the *Schrödinger suite (36)*. In the case of the gatekeeper mutants, the corresponding gatekeeper residues (tyrosine and threonine) were prepared in the same way.

The *def2TZVP* basis set was used for all calculations and effective core potentials (ECPs) were used for the iodine atom. *Ab initi*o interaction energies at the MP2 level were calculated using the *GAMESS* software, and partitioned into their constituent interaction energy terms (see Equation 1) using the many body interaction energy decomposition scheme (EDS) described by *Góra et al.*(37). In this scheme, the total interaction energy is calculated in a super-molecular approach as the difference between the total energy of a complex (here, of the inhibitor and the gatekeeper residue) and the sum of the energies of its isolated constituents. In all calculations, the complex centered basis set (CCBS) was used consistently and the results are therefore basis set superposition error (BSSE) free due to the full counterpoise correction.

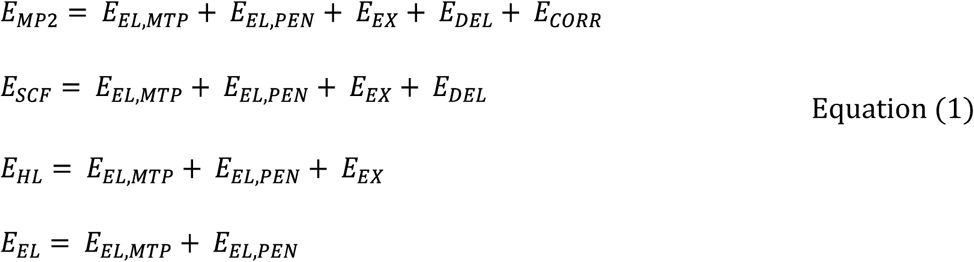

As shown in Equation 1, the total interaction energy at the MP2 level of theory (E_MP2_) includes the components of the Hartree-Fock interaction energy (*E*_*MP2*_) and the second order Coulomb correlation correction term (*E*_*CORR*_). This correlation term (*E*_*CORR*_) includes the second order intermolecular dispersion energy and the correlation corrections to the SCF components. The Hartree-Fock interaction energy (*E*_*SCF*_) was partitioned into a first order Heitler-London component (*E_HL_*) and a higher order Hartree-Fock delocalization interaction energy component (*E_DEL_*), which encompasses the induction and the associated exchange effects. Because their separation could lead to a non-physical charge transfer, this component was not partitioned any further. The Heitler-London interaction energy component (*E_HL_*) can be separated into the first-order electrostatic interactions (*E_EL_*) of monomers (the inhibitor and the gatekeeper residue in our case) and the associated Heitler-London exchange repulsion energy (*E_EX_*) due to the Fermi electron correlation effects. The electrostatic interaction energy (*E_EL_*) was obtained as a first-order term in the polarization perturbation theory and the exchange repulsion term (*E_EX_*) was calculated by subtracting the electrostatic interaction energy from the Heitler-London energy (*E_EX_* = *E_HL_* - *E_EL_*). *E_EL, MTP_* refers to the electrostatic multipole component estimated from an atomic multipole expansion, *E_EL,PEN_* is the electrostatic penetration energy, calculated from the following expression: *E_EL,PEN_* = *E_EL_* - *E_EL,MTP_*.

### Binding free energy calculations

The molecular mechanics-generalized Born surface area (MM/GBSA) method was used to estimate the binding free energy of the inhibitors to haspin kinase. The initial coordinates of the haspin-inhibitor complexes were obtained from the co-crystallized structures (see Supplementary Figure 2). The *Protein Preparation* wizard of the *Schrodinger suite* (*Version 2015.r3*) was used for pre-processing of the structures, formation of disulfide bonds, addition of hydrogen atoms and assigning protonation states at pH 7.0. The *pmemd* module of the *Amber14* software suite (38) was used to perform the molecular dynamics (MD) simulations with the *ff14SB*(39) force field for protein. The *LEap* module of *AmberTools14* was used to construct the topologies of the haspin-inhibitor complexes. The ligand parameters were generated based on the generalized Amber force field (*GAFF*). To improve the description of charge, dipole moment and geometry of halogenated compounds in molecular mechanics calculations, the positive region (σ hole) centered on the halogen atom was represented by an extra-point charge (EP). This inclusion of an EP results in improved modeling of halogen-bonding in MD simulations. The force field parameters for this EP were taken from *Ibrahim et al.(40)*. For generation of the partial atomic charges for the ligands, the *RESP(41)* program was used to fit the atom-centered charges to the molecular electrostatic potential (MEP) grid computed by the *GAMESS* program. The system was centred and aligned with the axes to minimize the volume. The system was then solvated using the *TIP3P* water model(42) by immersing the protein-ligand complex in a cubic box of water molecules, such that the shortest distance between the edge of the solvation box and the complex is 10 Å. The net charge (−2e) of the system was then neutralized by adding Na^+^ counter ions. For each system, energy minimization was performed in three 1500-cycle consecutive runs using the steepest descent minimization method followed by switching to the conjugate gradient method after 500 cycles. Gradually decreasing harmonic restraints with force constants of 500, 1 and 0 kcal/mol.Å^2^ were used for non-hydrogen atoms in three consecutive runs. Energy minimization was followed by 1 nanosecond (*ns*) of gradual heating from 10 K to 300 K with harmonic restraints with a force constant of 50 kcal/mol.Å^2^ acting on non-hydrogen atoms. Then the system was equilibrated for 1 *ns* under *NPT* conditions at 300K, with heavy atoms (except solvent ions) harmonically restrained with a force constant of 50 kcal/mol.Å^2^. This was followed by an *NPT* equilibration of 2 *ns* without any positional restraints. The potential energy function and atomic co-ordinates were calculated using a 2 femtoseconds (*fs*) time step. The *SHAKE*(43) algorithm was used to constrain all the bonds involving hydrogen atoms. The *Particle Mesh Ewald* (PME) method was used to calculate the electrostatic interactions. A cut-off of 10 Å was set for generating the non-bonded pair list and this pair list was updated every 100 steps. After equilibration, data were collected over a 6 *ns* simulation run for binding free energy calculations and 3000 sets of atomic coordinates were saved every 2 picoseconds (*ps*). MM/GBSA calculations of the binding free energy were performed using the *MMPBSA.py* module implemented in the *Amber14* analysis tools. A single-trajectory approach was used in which receptor, ligand and complex geometries were extracted from a single MD trajectory. All the ions and water molecules were stripped from the trajectory snapshots. A salt concentration of *0.15 M* and the Born implicit solvent model (*igb = 2*) were used. Each binding free energy was computed as the sum of a molecular mechanics term (ΔE_gas_), a Gibbs solvation term (Δ Δ G_solvation_) and an entropic contribution (TΔ S_solute_). For the entropic contribution to binding free energy, we computed translational and rotational entropies with a rigid rotor model using the *MMPBSA.py* module. The calculation of vibrational entropies using normal-mode analysis with MMPBSA.py failed due to the inclusion of the EP in the force field. The free energy of binding for some of the derivatives is positive since vibrational and conformational entropy terms are neglected.

**Supplemental Figure S1.**
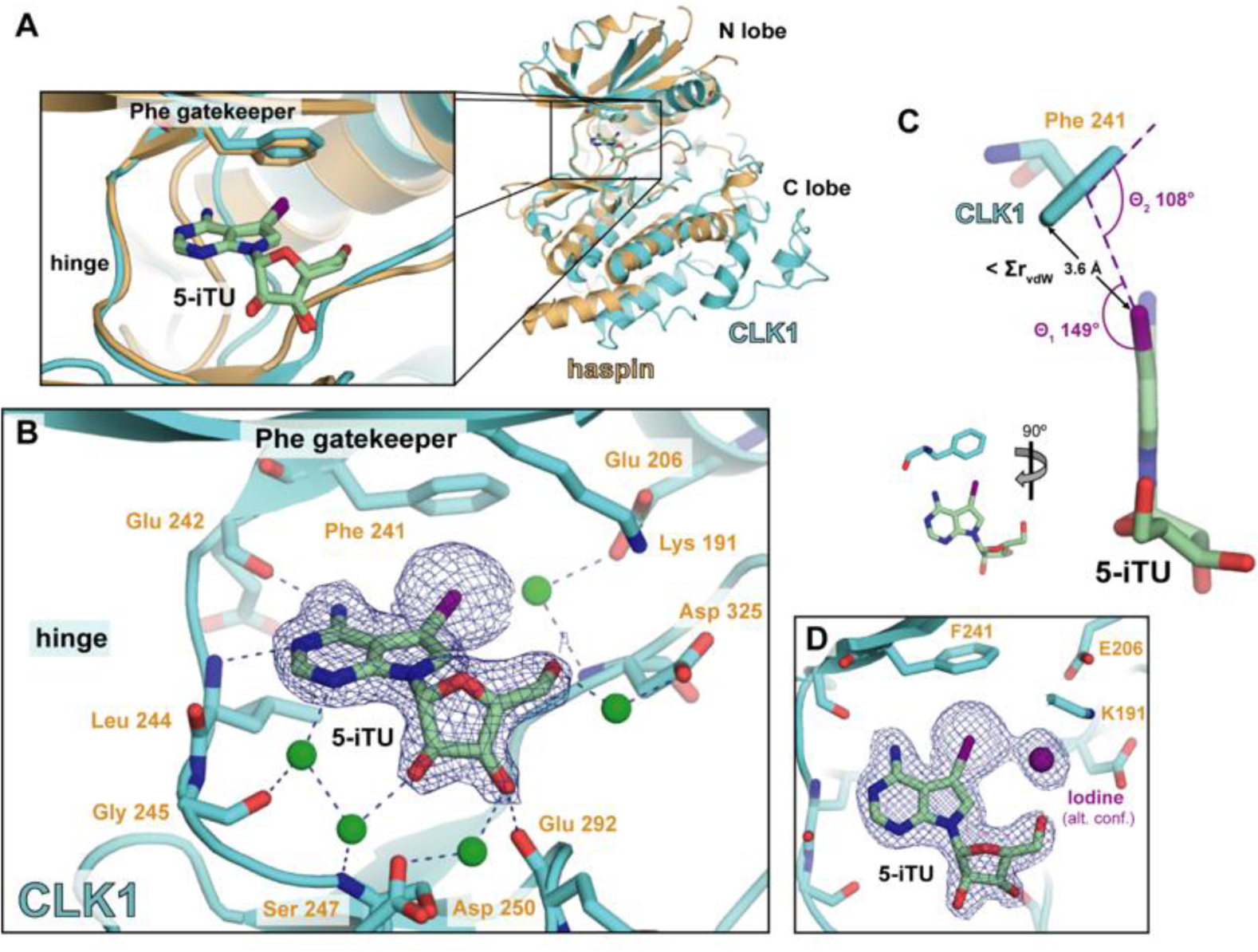
The binding mode of 5-iTU is maintained in CLK1. (**A**) Superimposition of the structures of CLK1 and haspin (PDB ID: 4OUC) in complex with 5-iTU, showing that the binding mode is conserved. (**B**) Co-crystal structure of CLK1 and 5-iTU, highlighting interacting amino acid residues and the Phe gatekeeper. Water molecules are represented as green spheres, hydrogen bonds within 3 Å as blue dashed lines. Relevant atoms are colored as follows: oxygen – red, nitrogen – blue and iodine – purple. |2Fo|-|Fc| omitted electron density map is contoured at 3σ level. (**C**) Geometric measures of the putative π-X bond. (**D**) |2Fo|-|Fc| omitted electron density map contoured at 1σ level, showing alternative conformation of delocalized iodine in binding pocket.

**Supplemental Figure S2.**
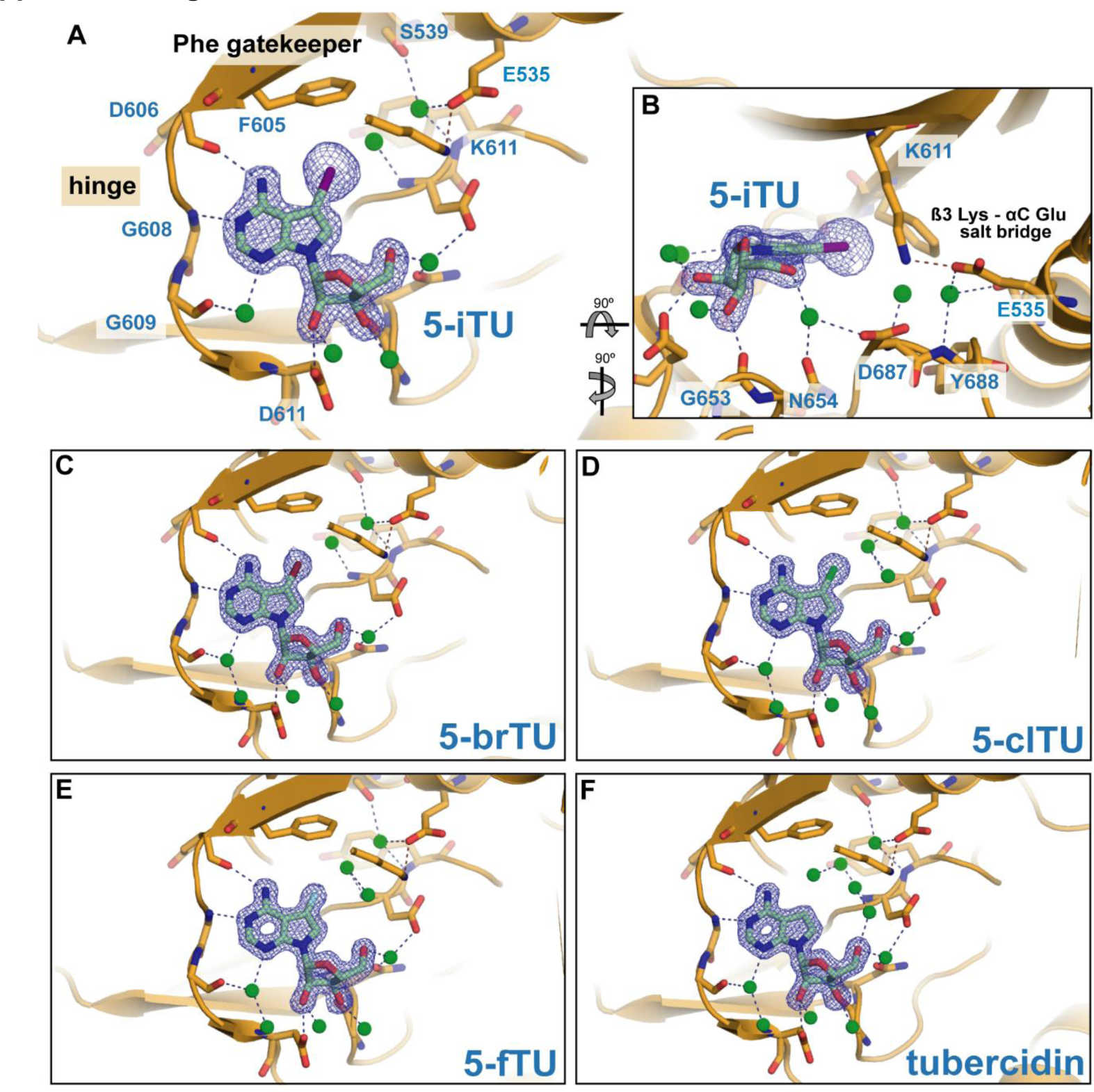
Structural studies of haspin in complex with the halogenated derivatives. (**A-B**) Co-crystal structure of haspin and 5-iTU, showing the ATP-competitive binding mode for the inhibitor. Interacting residues and the Phe gatekeeper are highlighted. Water molecules are represented as green spheres, hydrogen bonds within 3 Å as blue dashed lines. Relevant atoms are colored as follows: oxygen – red, nitrogen – blue and iodine – purple. |2Fo|-|Fc| omitted electron density map is contoured at 3σ level. (**C-F**) Co-crystal structures of haspin and different derivatives (see blue label), representation equivalent to (A). Bromine atom is shown in dark red, chlorine atom in green and fluorine atom in cyan.

**Supplemental Figure S3.**
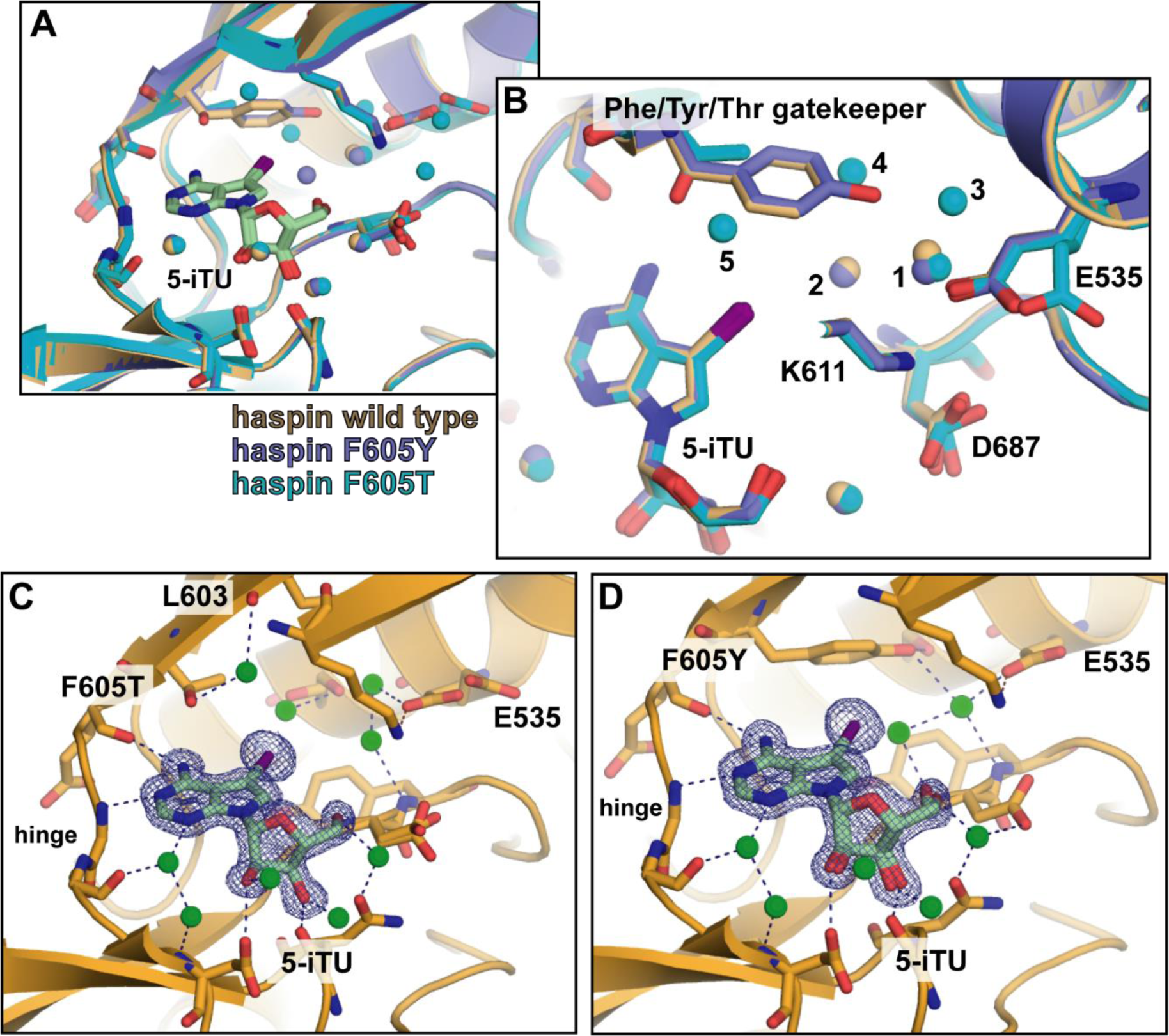
5-iTU-haspin binding site is maintained in the absence of an aromatic gatekeeper residue. (**A**) Superimposition of co-crystal structures of wild-type haspin (orange), haspin^F605Y^ (slate) and haspin^F605T^ (cyan) in complex with 5-iTU, showing that the binding modus is maintained in gatekeeper mutants. (**B**) Similar to (A), highlighting relevant residues in the binding pocket. Water molecules are shown as spheres and numbered from 1-5. (**C**) Co-crystal structure of haspin^F605T^ in complex with 5-iTU. Water molecules are represented as green spheres and hydrogen bonds within 3 Å as blue dashed lines. Relevant atoms are coloured as follows: oxygen – red, nitrogen – blue and iodine – purple. |2Fo|-|Fc| omitted electron density map is contoured at 3σ level. (**D**) Same as (C), showing the co-crystal structure of haspin^F605Y^ in complex with 5-iTU.

**Supplemental Figure S4.**
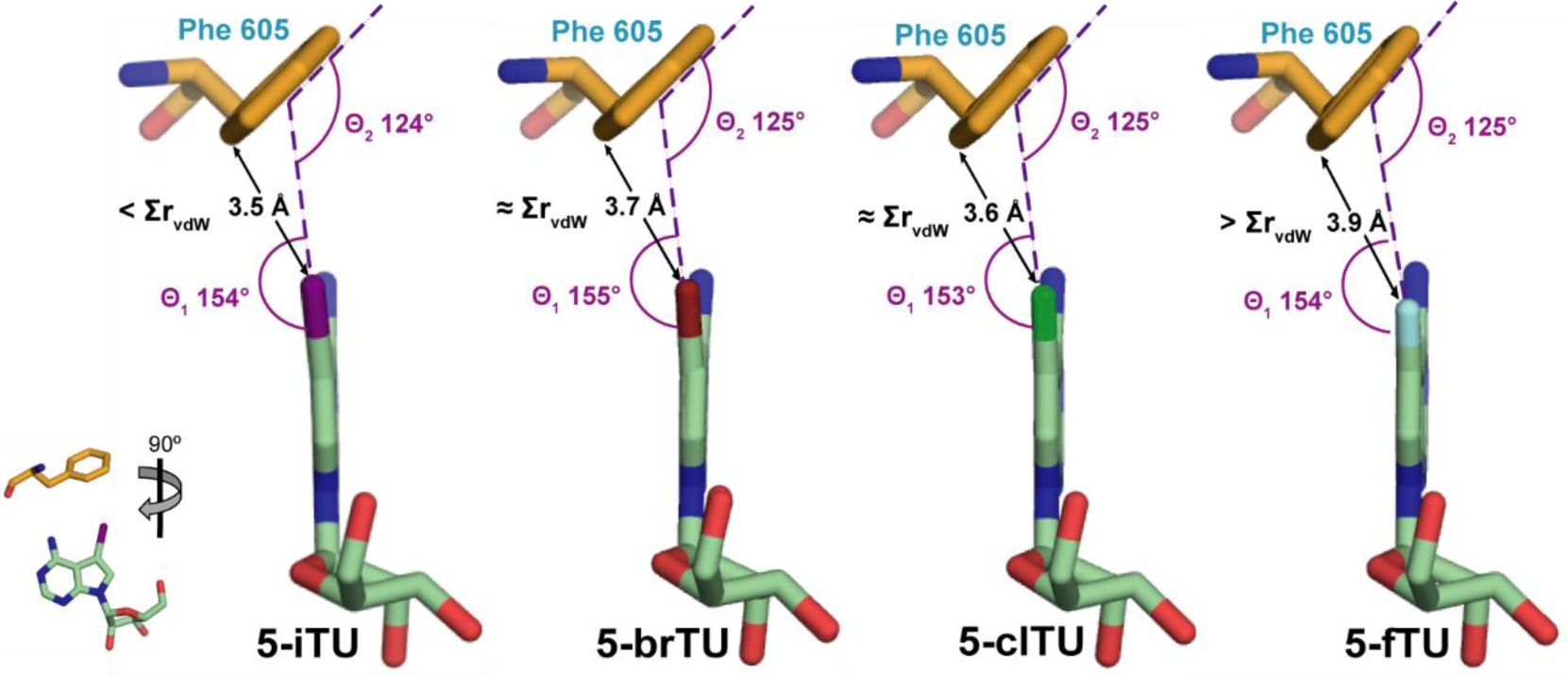
Geometric measures of putative halogen-π-bonds. Overview of the halogenated derivatives approaching the aromatic ring of the Phe gatekeeper residue in haspin. The distance between the halogen and the closest carbon atom of the aromatic ring was measured and compared with the sum of the van der Waals radii (∑r_vdW_). Θ1 is the angle between C-X bond to the centre of the phenylalanine aromatic group (C-X···π), while Θ_2_ is the angle of the halogen to the plane of phenylalanine aromatic ring (X⋯π-C).

**Supplemental Figure S5.**
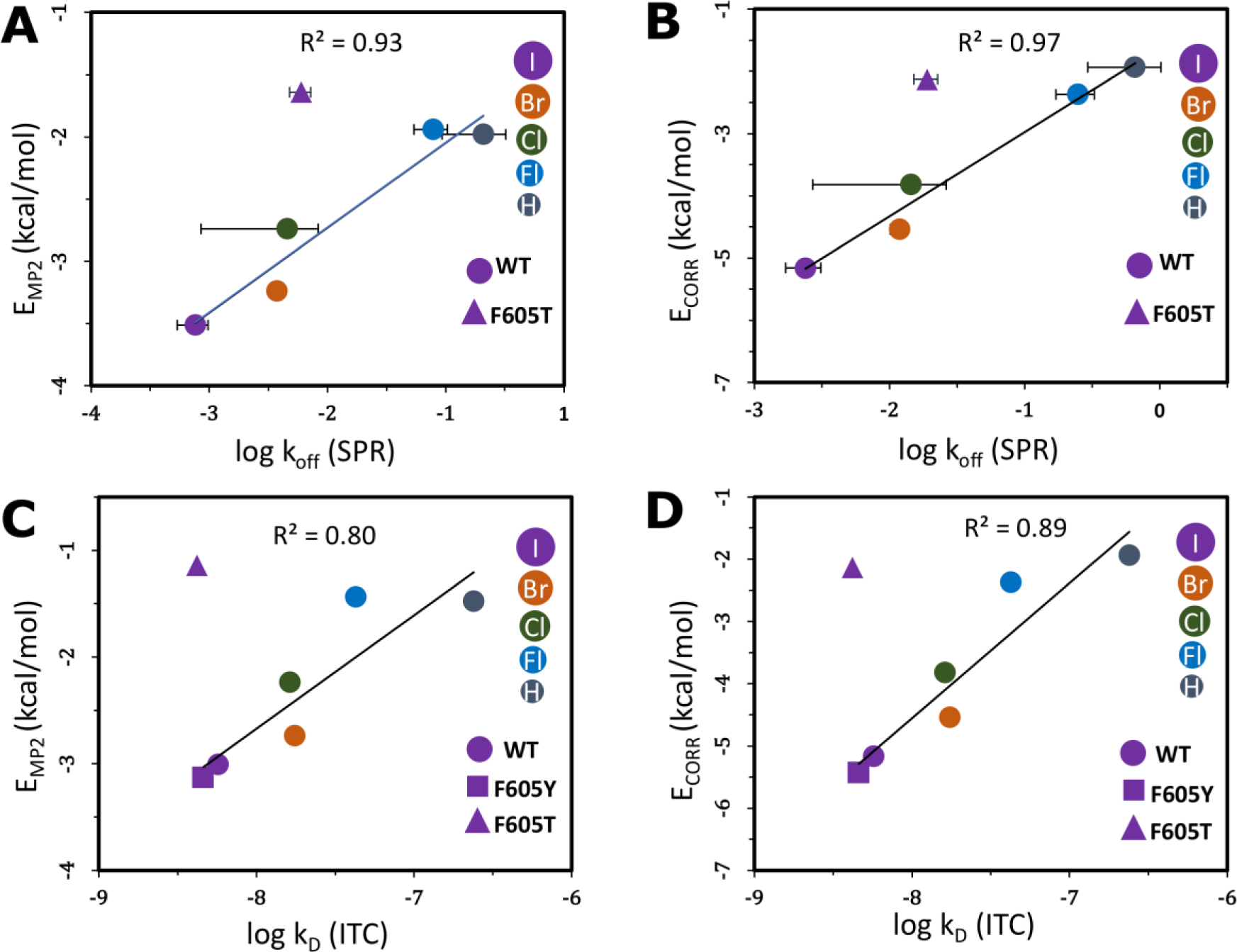
Correlation plots of computed quantum mechanical energies against experimental binding parameters. A) Second-order Møller–Plesset interaction energy (E_MP2_) between the tubercidin derivatives and the gatekeeper residue versus the experimental (SPR) dissociation rate constants (k_off_) of the tubercidin derivatives. B) The second-order correlation correction energy term (E_CORR_) for the interaction between the tubercidin derivatives and the gatekeeper residue versus the experimental (SPR) dissociation rate constants (k_off_) of the tubercidin derivatives This correlation energy (E_CORR_) includes second-order intermolecular dispersion interactions and the correlation corrections to the Hartree-Fock (HF) energy. C) Second-order Møller–Plesset interaction energy (E_MP2_) between tubercidin derivatives and gatekeeper residue versus the experimental (ITC) binding affinities (k_D_) of the tubercidin derivatives. D) Second-order correlation correction energy term (E_CORR_) for the interaction between the tubercidin derivatives and the gatekeeper residue versus the experimental (ITC) binding affinities (k_D_) of the tubercidin derivatives. The correlation coefficients (R^2^) and the linear fits were computed omitting the outlier data points for the F605T mutant. The error bars for the K_D_ (ITC) values are smaller than the size of the data point symbols.

**Supplemental Table S1.** DSF results for 5-iTU against 137 kinases. All results are listed with decreasing melting temperature shifts (ΔT_m_). The gatekeeper residues (GK) of the kinases were determined by sequence alignment and analysis of the structures for hits with ΔT_m_ > 2º.

| Kinase | $\Delta T_m$ | GK | Kinase | $\Delta T_m$ | GK | Kinase | $\Delta T_m$ | Kinase | $\Delta T_m$ |
| --- | --- | --- | --- | --- | --- | --- | --- | --- | --- |
| Haspin | 11.40 | Phe | TTK | 2.20 | Met | MAPK2TG | 0.79 | CAMK1G | 0.20 |
| DYRK2 | 10.38 | Phe | RSK4 | 2.20 | Leu | GRK1 | 0.76 | CAMK1D | 0.20 |
| CLK4 | 9.59 | Phe | AMPK1 | 2.19 | Met | MYLK | 0.73 | CDPK1PF | 0.12 |
| CLK1 | 8.60 | Phe | PIM2 | 2.04 | Leu | CHEK2 | 0.70 | VRK3 | 0.10 |
| CLK3 | 7.50 | Phe | GSK3β | 2.00 | Leu | CAMK2A | 0.70 | VRK2 | 0.10 |
| CK1ε | 5.90 | Met | CK2α2 | 1.97 | Phe | RIPK2 | 0.69 | TOPK | 0.10 |
| DYRK1A | 5.80 | Phe | MAP2K2 | 1.90 |  | MAPK9A | 0.60 | CAMK2B | 0.10 |
| CLK2 | 5.20 | Phe | PRKCL1 | 1.83 |  | CAMK2G | 0.60 | AAK1 | 0.09 |
| CDK2 | 4.98 | Phe | ITK | 1.78 |  | BMP2K | 0.60 | ADRBK2 | 0.03 |
| MST3 | 4.96 | Met | STK38 | 1.70 |  | RPS6KA1 | 0.50 | BMX | 0.02 |
| DRAK2 | 4.84 | Leu | SNF1LK | 1.70 |  | AMPKA2 | 0.50 | PRKG2 | 0.00 |
| LOK | 4.70 | Ile | RPS6KA2 | 1.70 |  | LOC340156 | 0.48 | PDK1 | 0.00 |
| YSK1 | 4.30 | Met | RIOK2 | 1.70 |  | ADRBK1 | 0.48 | NLK | -0.10 |
| SLK | 4.20 | Ile | PRKCL2 | 1.70 |  | MAPK13 | 0.46 | MAP2K6 | -0.10 |
| CAMK4 | 3.75 | Leu | MERTK | 1.70 |  | TEC | 0.45 | NEK6 | -0.20 |
| AMPKα2 | 3.58 | Met | PDPK1 | 1.69 | | CDKL2 | 0.43 | CDK6 | -0.20 |
| MPSK1 | 3.40 | Leu | RPS6KA3 | 1.66 |  | NEK7 | 0.40 | PKMYT1 | -0.22 |
| CK1γ2 | 3.30 | Leu | PRKCZ | 1.56 | | NEK2 | 0.40 | DDR1 | -0.38 |
| PHKγ2 | 3.24 | Phe | MAPK3A | 1.50 | | MAP3K5 | 0.40 | CDC42BPG | -0.38 |
| RIPK3 | 3.21 | Thr | TLK1 | 1.45 |  | JAK1 | 0.40 | MAPK11 | -0.40 |
| CDKL5 | 3.16 | Phe | PIM1 | 1.40 |  | SRPK2 | 0.39 | CAMK2D | -0.40 |
| MST1 | 3.10 | Met | NEK11 | 1.31 |  | CDPK1PV | 0.39 | DMPK1 | -0.58 |
| STK33 | 3.06 | Leu | PRKACA | 1.30 |  | BRK1 | 0.39 | CDC42BPAB | -0.74 |
| PLK4 | 2.90 | Leu | PIM3 | 1.30 |  | MAPK2PF | 0.35 |  |  |
| DAPK3 | 2.90 | Leu | GAK | 1.30 |  | CDC42BPB | 0.35 |  |  |
| DRAK1 | 2.80 | Leu | CDKL3 | 1.30 |  | ZAK | 0.34 |  |  |
| PFTAIRE1 | 2.71 | Phe | MYLK2 | 1.28 |  | DCAMKL | 0.33 |  |  |
| CDK8 | 2.71 | Phe | IKBKB | 1.25 |  | YANK3 | 0.30 |  |  |
| MST2 | 2.70 | Met | PRKD2 | 1.24 |  | PRKX | 0.28 |  |  |
| ERK3 | 2.70 | Gln | PCTK1 | 1.11 |  | TYK2 | 0.27 |  |  |
| CSNK1γ3 | 2.70 | Leu | NEK9 | 1.06 | | MSSK1 | 0.26 | | |
| PRKCN | 2.62 | Met | CSNK2A1 | 1.05 |  | LIMK1 | 0.26 |  |  |
| NDR2 | 2.60 | Met | TNIK | 1.00 |  | SRPK1 | 0.25 |  |  |
| CK1γ1 | 2.50 | Leu | PRKG1 | 1.00 | | VRK1 | 0.20 | | |
| QIK | 2.41 | Ile | NEK1 | 0.96 |  | TYRO3 | 0.20 |  |  |
| STK39 | 2.40 | Met | PCTK2 | 0.83 |  | PAK6 | 0.20 |  |  |
| MEK1 | 2.29 | Met | MST4 | 0.80 |  | PAK5 | 0.20 |  |  |
| GRK5 | 2.22 | Leu | YANK1 | 0.79 |  | CDKL1 | 0.20 |  |  |

**Supplemental Table S2A.** Supplemental Table 2. BLI and ITC data. Summary of ITC and BLI data for 5-iTU in complex with CLK1 and CLK3

| 5-iTU vs | ITC data |  |  |  |  |  | BLI data |  |  |
| --- | --- | --- | --- | --- | --- | --- | --- | --- | --- |
| | n | $\Delta H$<br>kcal/mol | $T\Delta S$<br>kcal/mol | $\Delta G$<br>kcal/mol | K<br>M<sup>-1</sup> | K<sub>d</sub><br>nM | $k_{on}$<br>M<sup>-1</sup> s<sup>-1</sup> | $k_{off}$<br>s<sup>-1</sup> | Resi-<br>denceTime |
| CLK1 | 0.96 | -16.66<br>(± 0.09) | -5.94 | -10.72 | 1.39e+8<br>(± 1.99e+7) | **7.2**<br>(±1.0) | 10.9e+4<br>(±8.48e+3) | 3.14e-4<br>(±1.06e-5) | **53.1 min**<br>(±6.3 min) |
| CLK3 | 1.00 | -12.04<br>(± 0.05) | -2.44 | -9.60 | 1.91e+7<br>(± 1.28e+6) | **52.4**<br>(±3.5) | - | - | - |

**Supplemental Table S2B.** Summary of ITC and BLI data for halogenated derivatives against haspin

| haspin vs | ITC data |  |  |  |  |  | BLI data |  |  |
| --- | --- | --- | --- | --- | --- | --- | --- | --- | --- |
| | n | $\Delta H$<br>kcal/mol | $T\Delta S$<br>kcal/mol | $\Delta G$<br>kcal/mol | K<br>M<sup>-1</sup> | K<sub>d</sub><br>nM | $k_{on}$<br>M<sup>-1</sup> s<sup>-1</sup> | $k_{off}$<br>s<sup>-1</sup> | Resi-<br>denceTime |
| tubercidin | 0.95 | -16.95<br>(± 0.10) | -8.21 | -8.74 | 4.18e+6<br>(±1.74e+5) | **239.2**<br>±10.0) | 9.56e+4<br>(±1.45e+3) | 4.74e-2<br>(±6.72e-4) | **21.1 sec**<br>(±0.3 sec) |
| 5-FTu | 1.00 | -18.10<br>(± 0.07) | -8.39 | -9.71 | 2.34e+7<br>(±1.56e+6) | **42.7**<br>(±2.8) | 9.26e+4<br>(±9.71e+2) | 6.32e-3<br>(±4.44e-5) | **2.6 min**<br>(±1.1 sec) |
| 5-ClTu | 0.98 | -21.54<br>(± 0.06) | -11.27 | -10.27 | 6.14e+7<br>(±3.29e+6) | **16.3**<br>(±0.9) | 9.00e+4<br>(±5.94e+2) | 9.22e-4<br>(±8.00e-6) | **18.1 min**<br>(±9.4 sec) |
| 5-BrTu | 1.00 | -22.50<br>(±0.07) | -12.28 | -10.22 | 5.70e+7<br>(±4.26e+6) | **17.5**<br>(±1.3) | 9.57e+4<br>(±3.36e+2) | 6.42e-4<br>(±5.76e-5) | **25.9 min**<br>(±2.3 min) |
| 5-ITu | 0.96 | -23.40<br>(±0.07) | -12.53 | -10.87 | 1.75e+8<br>(±1.66e+7) | **5.7**<br>(±0.5) | 9.59e+4<br>(±5.61e+2) | 2.76e-4<br>(±5.96e-6) | **60.4 min**<br>(±1.3 min) |

**Supplemental Table S2C.** Summary of ITC and BLI data for haspin gatekeeper mutants against 5-iTU

| 5-iTU vs | ITC data |  |  |  |  |  | BLI data |  |  |
| --- | --- | --- | --- | --- | --- | --- | --- | --- | --- |
| | n | $\Delta H$<br>kcal/mol | $T\Delta S$<br>kcal/mol | $\Delta G$<br>kcal/mol | K<br>M<sup>-1</sup> | K<sub>d</sub><br>nM | $k_{on}$<br>M<sup>-1</sup> s<sup>-1</sup> | $k_{off}$<br>s<sup>-1</sup> | Resi-<br>denceTime |
| F605Y | 0.96 | -22.26<br>(± 0.05) | -11.27 | -10.99 | 2.34e+7<br>(± 1.56e+6) | **4.6**<br>(±0.3) | 11.2e+4<br>(±1.45e+3) | 4.29e-4<br>(±8.87e-6) | **38.9 min**<br>(±48 sec) |
| F605H | 0.98 | -20.35<br>(± 0.05) | -9.42 | -10.93 | 6.14e+7<br>(± 3.29e+6) | **5.1**<br>(±0.3) | - | - | - |
| F605M | 0.94 | -22.09<br>(±0.04) | -10.89 | -11.20 | 5.70e+7<br>(± 4.26e+6) | **3.2**<br>(±0.2) | - | - | - |
| F605L | 0.97 | -23.35<br>(±0.04) | -12.36 | -10.99 | 1.75e+8<br>(± 1.66e+7) | **4.6**<br>(±0.4) | - | - | - |
| F605T | 0.93 | -20.48<br>(±0.03) | -9.42 | -11.06 | 1.75e+8<br>(± 1.66e+7) | **4.2**<br>(±0.4) | 12.4e+4<br>(±6.22e+2) | 1.23e-3<br>(±6.54e-6) | **13.5 min**<br>(±4.3 sec) |
| F605Q | 0.98 | -23.23<br>(±0.04) | -12.56 | -10.67 | 1.75e+8<br>(± 1.66e+7) | **8.0**<br>(±0.8) | - | - | - |

**Supplemental Table S3.** Summary of SPR data for haspin wild-type and F605T mutant against tubercidin derivatives

| | $K_D$ eq<br>(nM) | $K_D$ kin<br>(nM) | $k_{on}$<br>($M^{-1}s^{-1}$) | $k_{off}$<br>($s^{-1}$) |
| --- | --- | --- | --- | --- |
| tubercidin<br>(hydrogen) | 2560<br>$\pm 1230$ | 2910<br>$\pm 707$ | $1.96 \times 10^5$<br>$\pm 1.59 \times 10^5$ | $6.57 \times 10^{-1}$<br>$\pm 3.62 \times 10^{-1}$ |
| 5-ftU<br>(fluoride) | 155<br>$\pm 91$ | 79.3<br>$\pm 35.9$ | $2.06 \times 10^6$<br>$\pm 2.07 \times 10^6$ | $2.49 \times 10^{-1}$<br>$\pm 0.79 \times 10^{-1}$ |
| 5-clTU<br>(Chloride) | 6.6<br>$\pm 2.8$ | 4.9<br>$\pm 2.8$ | $2.79 \times 10^6$<br>$\pm 1.26 \times 10^6$ | $1.45 \times 10^{-2}$<br>$\pm 1.18 \times 10^{-2}$ |
| 5-brTU<br>(Bromide) | 2.6<br>$\pm 0.3$ | 1.3<br>$\pm 0.3$ | $9.71 \times 10^6$<br>$\pm 2.67 \times 10^6$ | $1.19 \times 10^{-2}$<br>$\pm 0.18 \times 10^{-2}$ |
| 5-iTU<br>(iodide) | 0.8<br>$\pm 0.2$ | 0.3<br>$\pm 0.1$ | $9.39 \times 10^6$<br>$\pm 2.03 \times 10^6$ | $2.40 \times 10^{-3}$<br>$\pm 0.70 \times 10^{-3}$ |
| fold difference |  |  |  |  |
| H to I | ↓ 3200 | ↓ 9700 | ↑ 48 | ↓ 274 |
| F to I | ↓ 194 | ↓ 264 | ↑ 5 | ↓ 104 |

**B** mutant F605T
| | $K_D$ eq<br>(nM) | $K_D$ kin<br>(nM) | $k_{on}$<br>($M^{-1}s^{-1}$) | $k_{off}$<br>($s^{-1}$) |
| --- | --- | --- | --- | --- |
| tubercidin<br>(hydrogen) | 5410<br>$\pm 2180$ | 6870 | $5.53 \times 10^4$<br>$\pm 1.25 \times 10^4$ | $3.29 \times 10^{-1}$<br>$\pm 0.68 \times 10^{-1}$ |
| 5-ftU<br>(fluoride) | 1660<br>$\pm 661$ | 1490<br>$\pm 258$ | $1.22 \times 10^5$<br>$\pm 0.40 \times 10^5$ | $1.76 \times 10^{-1}$<br>$\pm 0.33 \times 10^{-1}$ |
| 5-clTU<br>(Chloride) | 166<br>$\pm 81$ | 154<br>$\pm 44$ | $7.47 \times 10^5$<br>$\pm 1.00 \times 10^5$ | $1.12 \times 10^{-1}$<br>$\pm 0.25 \times 10^{-1}$ |
| 5-brTU<br>(Bromide) | 53.3<br>$\pm 24.6$ | 46.3<br>$\pm 9.1$ | $1.07 \times 10^6$<br>$\pm 0.12 \times 10^6$ | $4.88 \times 10^{-2}$<br>$\pm 0.77 \times 10^{-2}$ |
| 5-iTU<br>(iodide) | 12.8<br>$\pm 5.1$ | 12.0<br>$\pm 2.2$ | $1.61 \times 10^6$<br>$\pm 0.44 \times 10^6$ | $1.89 \times 10^{-2}$<br>$\pm 0.38 \times 10^{-2}$ |
| fold difference |  |  |  |  |
| H to I | ↓ 422 | ↓ 573 | ↑ 29 | ↓ 17 |
| F to I | ↓ 129 | ↓ 124 | ↑ 13 | ↓ 9 |

| fold difference |  |  |  |  |
| --- | --- | --- | --- | --- |
| | $K_D$ eq<br>(nM) | $K_D$ kin<br>(nM) | $k_{on}$<br>($M^{-1}s^{-1}$) | $k_{off}$<br>($s^{-1}$) |
| 5-iTU | ↓ 16 | ↓ 40 | ↑ 6 | ↓ 8 |

**Supplemental Table S4.** Solvent accessible surface area of ordered water molecules in the binding pocket. The Solvent Accessible Surface Area (SASA) of water molecules W1-W5 (as shown in Supplemental Figure 3), the nitrogen atom of the ε-amino group of K511 (NZ) and the closest C atom of the aromatic ring of F605 (CD1), was obtained by the POPS server, using a probe with 1.4 Å radius(44). The structures with the halogenated derivatives are compared with the haspin apo-structure (PDB ID: 2WB8).

| SASA (Å<sup>2</sup>) | haspin/<br>5-iTU | haspin/<br>5-brTU | haspin/<br>5-clTU | haspin/<br>5-ftTU | haspin/<br>tubercidin | Apo haspin |
| --- | --- | --- | --- | --- | --- | --- |
| W1 | **2.13** | **2.20** | **1.88** | **1.84** | **1.74** | **1.34** |
| W2 | **3.14** | **2.69** | **2.64** | **2.64** | **1.70** | **2.06** |
| W3 | - | - | 4.03 | 3.94 | 3.50 | 5.07 |
| W4 | - | - | - | - | 3.47 | - |
| W5 | - | - | - | - | 3.21 | - |
| NZ (K511) | **4.56** | **4.88** | **3.80** | **3.46** | **4.02** | **4.23** |
| CD<sub>1</sub> (F605) | **1.49** | **1.39** | **1.44** | **1.41** | **1.37** | **1.31** |

**Supplemental Table S5.** Measured geometric parameters between the halogens and the Phe gatekeeper

|  | CLK1 /<br>5-iTU | haspin /<br>5-iTU | haspin /<br>5-brTU | haspin /<br>5-clTU | haspin /<br>5-ftTU | Preferred ge-<br>ometry of<br>X-bonds |
| --- | --- | --- | --- | --- | --- | --- |
| Distance between<br>halogen and<br>closest C atom of<br>gatekeeper | **3.58 Å**<br>$\pm 0.28$ Å<br>($< \sum r_{vdW}$) | **3.52 Å**<br>$\pm 0.18$ Å<br>($< \sum r_{vdW}$) | **3.68 Å**<br>$\pm 0.19$ Å<br>($\approx \sum r_{vdW}$) | **3.61 Å**<br>$\pm 0.17$ Å<br>($\approx \sum r_{vdW}$) | **3.85 Å**<br>$\pm 0.17$ Å<br>($> \sum r_{vdW}$) | $\leq \sum r_{vdW}$ |
| Sum of van der<br>Waals radii | 3.68 Å | 3.68 Å | 3.60 Å | 3.45 Å | 3.20 Å | - |
| Angle $\Theta_1$ | 149.5° | 153.7° | 154.7° | 153.3° | 154.0° | $\approx 160^\circ - 165^\circ$ |
| Angle $\Theta_2$ | 108.0° | 124.0° | 125.1° | 124.6° | 125.3° | $\approx 120^\circ$ ($\approx 90^\circ$<br>for π-x-bond) |

**Supplemental Table S6.** Total interaction energy [*kcal. mol^-1^*] between tubercidin derivatives and gatekeeper Phe 605 residue at consecutively increasing levels of quantum mechanical theory, see equation 1. *E_EL_* is the electrostatic energy only, *E_HL_* includes the Heitler-London energy, *E_SCF_* includes the Hartree-Fock energy as well, and *E_MP2_* is the full Moeller-Plesset second order energy. k_off_ values were measured by SPR and BLI, and KD values were measured by SPR and ITC.

| Inhibitor | log $k_{\text{off}}$<br>(SPR) | log $k_{\text{off}}$<br>(BLI) | log $K_D$<br>(SPR) | log $K_D$<br>(ITC) | $E_{EL}$ | $E_{HL}$ | $E_{SCF}$ | $E_{MP2}$ |
| --- | --- | --- | --- | --- | --- | --- | --- | --- |
| 5-iTU | -2.62 | -3.56 | -9.10 | -8.24 | -1.85 | 2.79 | 2.15 | -3.01 |
| 5-brTU | -1.92 | -3.19 | -8.59 | -7.76 | -1.09 | 2.22 | 1.78 | -2.74 |
| 5-clTU | -1.84 | -3.04 | -8.18 | -7.79 | -0.52 | 1.91 | 1.58 | -2.24 |
| 5-ftTU | -0.60 | -2.20 | -6.81 | -7.37 | 0.17 | 1.15 | 0.94 | -1.44 |
| tubercidin | -0.18 | -1.32 | -5.59 | -6.62 | -0.09 | 0.65 | 0.47 | -1.48 |
| Pearson Correlation coefficient ($R^2$) with log $k_{\text{off}}$ (SPR) | | | | | 0.82 | -0.98 | -0.98 | 0.93 |
| Pearson Correlation coefficient ($R^2$) with log $k_{\text{off}}$ (BLI) | | | | | 0.82 | -0.98 | -0.99 | 0.91 |
| Pearson Correlation coefficient ($R^2$) with log $K_D$ (SPR) | | | | | 0.71 | -0.97 | -0.99 | 0.87 |
| Pearson Correlation coefficient ($R^2$) with log $K_D$ (ITC) | | | | | 0.79 | -0.96 | -0.97 | 0.86 |

**Supplemental Table S7.** Contribution of the different interaction energy terms to the total interaction energy, *E_MP2_* [*kcal. mol*^-1^], between tubercidin derivatives and the gatekeeper Phe 605 residue. See equation 1 for the definition of the terms. *E_EL,MTP_* is the electrostatic multipole term, *E_EL,PEN_* is the penetration electrostatic term, *E_EX_* is the exchange term, *E_DEL_* is the delocalization term, and *E_CORR_* is the correlation energy term. k_off_ values were measured by SPR and BLI, and KD values were measured by SPR and ITC.

| Inhibitor | log $k_{\text{off}}$<br>(SPR) | log $k_{\text{off}}$<br>(BLI) | log $K_D$<br>(SPR) | log $K_D$<br>(ITC) | $E_{EL,MTP}$ | $E_{EL,PEN}$ | $E_{EX}$ | $E_{DEL}$ | $E_{CORR}$ |
| --- | --- | --- | --- | --- | --- | --- | --- | --- | --- |
| 5-iTU | -2.62 | -3.56 | -9.10 | -8.24 | 2.48 | -4.33 | 4.63 | -0.64 | -5.16 |
| 5-brTU | -1.92 | -3.19 | -8.59 | -7.76 | 1.03 | -2.12 | 3.30 | -0.42 | -4.54 |
| 5-clTU | -1.84 | -3.04 | -8.18 | -7.79 | -0.21 | -0.32 | 2.44 | -0.34 | -3.82 |
| 5-ftTU | -0.60 | -2.20 | -6.81 | -7.37 | 0.13 | 0.04 | 0.97 | -0.21 | -2.37 |
| tubercidin | -0.18 | -1.32 | -5.59 | -6.62 | -0.07 | -0.01 | 0.74 | -0.18 | -1.94 |
| Pearson Correlation coefficient ($R^2$) with log $k_{\text{off}}$ (SPR) | | | | | -0.55 | 0.69 | -0.94 | 0.89 | 0.98 |
| Pearson Correlation coefficient ($R^2$) with log $k_{\text{off}}$ (BLI) | | | | | -0.68 | 0.76 | -0.92 | 0.88 | 0.96 |
| Pearson Correlation coefficient ($R^2$) with log $K_D$ (SPR) | | | | | -0.49 | 0.61 | -0.88 | 0.80 | 0.95 |
| Pearson Correlation coefficient ($R^2$) with log $K_D$ (ITC) | | | | | -0.70 | 0.76 | -0.90 | 0.88 | 0.92 |

**Supplemental Table S8.** Total interaction energy [*kcal. mol*^-1^] between tubercidin derivatives and gatekeeper Phe 605 residue at consecutively increasing levels of quantum mechanical theory, see equation 1. *E_EL_* is the electrostatic energy only, *E_HL_* includes the Heitler-London energy, *E_SCF_* includes the Hartree-Fock energy as well, and *E_MP2_* is the full Moeller-Plesset second order energy. k_off_ values were measured by SPR and BLI, and K_D_ values were measured by SPR and ITC. The correlation coefficient was calculated only for k_off_ and K_D_ values measured by BLI and ITC, respectively, as there were no SPR data available for the F605Y mutant.

| System | log $k_{\text{off}}$<br>(SPR) | log $k_{\text{off}}$<br>(BLI) | log $K_D$<br>(SPR) | log $K_D$<br>(ITC) | $E_{\text{EL}}$ | $E_{\text{HL}}$ | $E_{\text{SCF}}$ | $E_{\text{MP2}}$ |
| --- | --- | --- | --- | --- | --- | --- | --- | --- |
| Wild type | -2.62 | -3.56 | -9.10 | -8.24 | -1.85 | 2.79 | 2.15 | -3.01 |
| F605Y | ND | -3.37 | ND | -8.34 | -2.27 | 3.01 | 2.30 | -3.13 |
| F605T | -1.72 | -2.91 | -7.89 | -8.38 | -0.78 | 1.38 | 0.99 | -1.14 |
| Pearson Correlation coefficient<br>($R^2$) with log $k_{\text{off}}$ (BLI) | | | | | **0.84** | **-0.91** | **-0.92** | **0.94** |
| Pearson Correlation coefficient<br>($R^2$) with log $K_D$ (ITC) | | | | | **-0.52** | **0.64** | **0.65** | **-0.69** |

**Supplemental Table S9.** Contribution of the different interaction energy terms to the total interaction energy, *E_MP2_* [*kcal. mol^-1^*] between 5-iTU and the gatekeeper residue for the wild type and the two mutants. See equation 1 for the definition of the terms. *E_EL,MTP_* is the electrostatic multipole term, *E_EL,PEN_* is the penetration electrostatic term, *E_EX_* is the exchange term, *E_DEL_* is the delocalization term, and *E_CORR_* is the correlation energy term. k_off_ values were measured by SPR and BLI, and K_D_ values were measured by SPR and ITC. The correlation coefficient was calculated only for k_off_ and K_D_ values measured by BLI and ITC, respectively, as there were no SPR data available for the F605Y mutant.

| System | log $k_{\text{off}}$<br>(SPR) | log $k_{\text{off}}$<br>(BLI) | log $K_D$<br>(SPR) | log $K_D$<br>(ITC) | $E_{\text{EL,MTP}}$ | $E_{\text{EL,PEN}}$ | $E_{\text{EX}}$ | $E_{\text{DEL}}$ | $E_{\text{CORR}}$ |
| --- | --- | --- | --- | --- | --- | --- | --- | --- | --- |
| Wild type | -2.62 | -3.56 | -9.10 | -8.24 | 2.48 | -4.33 | 4.63 | -0.64 | -5.16 |
| F605Y | ND | -3.37 | ND | -8.34 | 0.41 | -2.68 | 5.28 | -0.71 | -5.43 |
| F605T | -1.72 | -2.91 | -7.89 | -8.38 | -6.97 | 6.19 | 2.17 | -0.39 | -2.13 |
| Pearson Correlation coefficient<br>($R^2$) with log $k_{\text{off}}$ (BLI) | | | | | **-1.00** | **0.99** | **-0.88** | **0.88** | **0.93** |
| Pearson Correlation coefficient<br>($R^2$) with log $K_D$ (ITC) | | | | | **0.86** | **-0.82** | **0.58** | **-0.57** | **-0.68** |

**Supplemental Table S10.** Binding free energies calculated using the MMGBSA approach for the binding of tubercidin derivatives with haspin.

| Inhibitor | $\Delta E_{\text{gas}} + \Delta \Delta G_{\text{solvation}}$<br>(Kcal/mol) | $T\Delta S_{\text{MMGBSA}}$<br>(Kcal/mol) | $\Delta G_{\text{MMGBSA}}$<br>(Kcal/mol) |
| --- | --- | --- | --- |
| 5-iTU | $-25.95 \pm 2.93$ | $-23.52 \pm 0.02$ | $-2.43 \pm 2.95$ |
| 5-brTU | $-26.00 \pm 2.75$ | $-23.27 \pm 0.02$ | $-2.73 \pm 2.77$ |
| 5-clTU | $-21.58 \pm 3.33$ | $-22.97 \pm 0.02$ | $1.39 \pm 3.35$ |
| 5-ftTU | $-20.31 \pm 3.04$ | $-22.83 \pm 0.01$ | $2.52 \pm 3.05$ |
| tubercidin | $-19.45 \pm 2.87$ | $-22.65 \pm 0.02$ | $3.20 \pm 2.89$ |

**Supplemental Table SS11.**
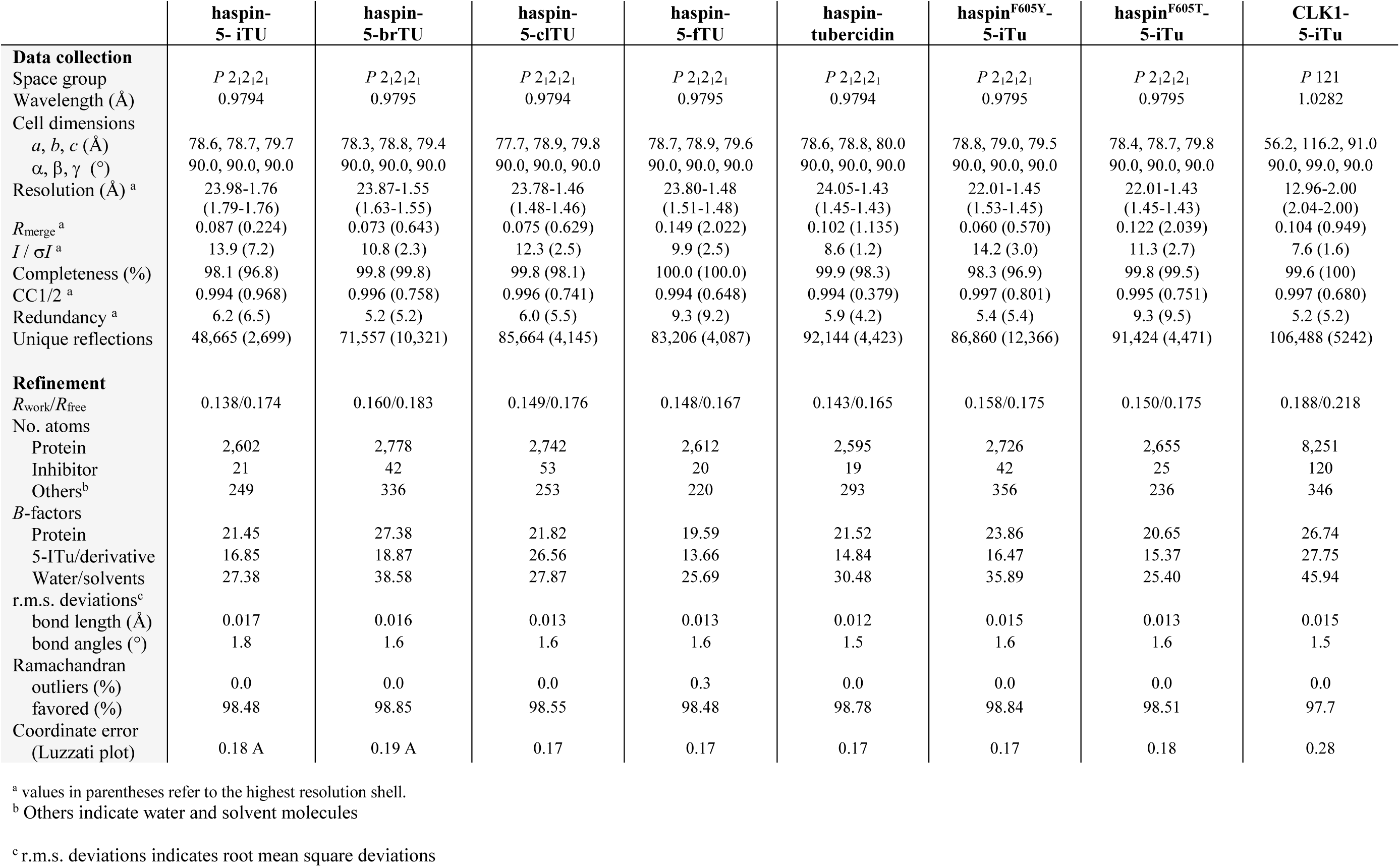
Data collection and refinement statistics

## Significance

The time that a drug resides on its target protein has emerged as an important parameter for drug development. A longer drug-target residence time can lead to improved drug efficacy and reduced adverse side effects. However, general strategies for the rational design of drugs with long target residence times are lacking. Here, by experimental and computational characterization of a set of inhibitor-kinase complexes, we show how inter-actions between inhibitor halogen atoms and protein aromatic residues can increase target residence times. Our results provide a simple strategy for the rational design of inhibitors with prolonged target residence times.

